# Evolution of co-regulatory network of C_4_ metabolic genes and TFs in the genus Flaveria: go anear or away in the intermediate species?

**DOI:** 10.1101/2020.10.02.324558

**Authors:** Ming-Ju Amy Lyu, Jemaa Essemine, Faming Chen, Genyun Chen, Xin-Guang Zhu

## Abstract

C_4_ photosynthesis evolved from the ancestral C_3_ photosynthesis by recruiting pre-existing genes to fulfill new functions. The enzymes and transporters required for the C_4_ photosynthesis have been intensively studied; however, the transcription factors (TFs) regulating these C_4_ metabolic genes are not well understood. In particular, how the TF regulatory network of C_4_ metabolic genes was rewired during the evolution is unclear. Here, we constructed TFs co-regulatory networks for core C_4_ metabolic genes (C_4_GRN) for four evolutionarily closely related species from the genus Flaveria, which represent four different evolutionary stages of the C_4_ photosynthesis, namely, C_3_, type I C_3_-C_4_, type II C_3_-C_4_ and C_4_. Our results show that more than half of the co-regulations of TFs and C_4_ core metabolic genes were species specific. The counterparts of C_4_ genes in C_3_ species were already co-regulated with the photosynthesis-related genes; whereas the required TFs for the C_4_ photosynthesis were recruited later. The type I C_3_-C_4_ species recruited 40% of C_4_ required TFs which co-regulated all core C_4_ metabolic genes but PEPC; nevertheless, the type II C_3_-C_4_ species took on a high divergent C_4_GRN with C_4_ species itself. In C_4_ species, PEPC and PPDK-RP possessed much more co-regulated TFs than other C_4_ metabolic genes. This study provides for the first time the TFs profiles of the C_4_ metabolic genes in species with different photosynthetic types and reveal the dynamic of C_4_ genes-TFs co-regulations along the evolutionary process, providing thereby new insights into the evolution of C_4_ photosynthesis.

## Introduction

C_4_ photosynthesis evolved from the ancestral C_3_ type (Sage 2004). In dual cell C_4_ species, C_4_ photosynthesis compartmentalizes the CO_2_ fixation in two cells, where CO_2_ is initially converted into a four-carbon acid by phospho*enol*pyruvate carboxylase (PEPC), which occurs mainly in the mesophyll cell (MC). The four-carbon acid diffuses then to nearby the bundle cell (BSC), where CO_2_ is subsequently released to be eventually fixed by the Calvin-Benson cycle (Hatch 1987). This effective synergistic collaboration between the two cells generates a CO_2_ concentrating mechanism (CCM), which results in a high CO_2_ concentration around the ribulose-1,5-bisphosphate carboxylase/oxygenase (Rubisco) in the BSC, decreasing thereby the photorespiration rate and leading ultimately to high light, water and nitrogen use efficiencies (Zhu, et al. 2008; Vogan and Sage 2011). The high C_4_ photosynthesis efficiency makes it an ideal target and promising to be engineered into C_3_ crops for the purpose of increasing the crop yield and productivity (Hibberd, et al. 2008; Sage and Zhu 2011).

All the involved enzymes and transporters in the C_4_ photosynthesis are recruited from pre-existing genes of the ancestral C_3_ species (Christin, et al. 2013; Moreno-Villena, et al. 2018). So far, the genes encoding the enzymes and transporters in the C_4_ core metabolism are well studied (Hatch 1987). The evolution of the C_4_ genes and transporters has also been well documented (Christin, et al. 2009; Williams, et al. 2012; Christin, et al. 2013; Moreno-Villena, et al. 2018). However, the regulatory mechanism controlling the C_4_ genes remains largely unknown (Hibberd and Covshoff 2010; Schluter and Weber 2020) and also its (regulatory mechanism) evolution constitutes a puzzling enigma. A recent study comparing for the first time the genome-wide transcriptional factor binding sites in three grass C_4_ and one grass C_3_ species provides a TF binding repertoire in C_4_ and C_3_ grass species, from which a landscape of TF families associated with those motifs are uncovered (Burgess, et al. 2019). Nevertheless, certain TFs regulating the C_4_ genes, especially the regulatory network regrouping the C_4_ genes and TFs co-regulations still remains poorly understood (Schluter and Weber 2020). It has been well documented that the *cis*-elements of C_4_ genes exist in their orthologs in C_3_ plants (Brown, et al. 2011; Kajala, et al. 2012; Burgess, et al. 2016); moreover, the *cis*-elements of C_4_ genes controlling the cell specificity could be recognized by the TFs in the C_3_ species and result in same cell specificity as C_4_ species (Brown, et al. 2011; Gorska, et al. 2019; Gupta, et al. 2020). However, it remains unclear to what extent the regulation of C_4_ genes required for C_4_ photosynthesis could be present in the C_3_ species, especially in intermediate species.

Among the model species used to study the C_4_ evolution, the genus Flaveria stands out since it comprises species with different photosynthetic types (Powell 1978; McKown, et al. 2005), in which the C_4_ species had evolved within 5 million years ago (Christin, et al. 2011). So far, the metabolic and anatomical features of C_3_, type I C_3_-C_4_, type II C_3_-C_4_, C_4_-like and C_4_ species in this genus have been extensively studied during the last decade (Sage 2004; McKown and Dengler 2007; Sage, et al. 2012; Mallmann, et al. 2014). The emergence of the type I C_3_-C_4_ is considered as prerequisite for the C_4_ evolution since the formation of C_2_ CCM (also known as photorespiratory carbon pump) creates an obvious input of amino residues from MC into BSC, which requires the rewiring of many C_4_ photosynthesis genes (Mallmann, et al. 2014). The type II C_3_-C_4_ species gained an increased C_4_-ness features, *e.g*., about 42% CO_2_ was initially fixed by PEPC (Rumpho, et al. 1984). It is worthy to note that both C_4_-like and C_4_ species have fully functional C_4_ metabolic pathways with the former (C_4_-like) still has small fraction of CO_2_ being initially fixed by Rubisco (Cheng, et al. 1988; Dmoore, et al. 1989). Thus, it represents an optimization from the C_4_-like to the C_4_ stage (Sage, et al. 2012). Though, an extensive knowledge has been accumulated on the sequences of metabolic changes in the different Flaveria species during the evolution of C_4_ photosynthesis, the knowledge about the molecular mechanisms underlying these changes was limited and was gained only based on a limited number of genes such as carbonic anhydrase (CA) (Gowik, et al. 2017), PEPC (Gowik, et al. 2004; Akyildiz, et al. 2007), and PEPC kinase (PEPC-k) (Aldous, et al. 2014).

The changes in the genetic regulatory mechanisms usually underlie the progressive changes in the adaptation and evolution of biological traits (Thompson, et al. 2015). Therefore, elucidation of the changes in genetic regulatory mechanisms during C_4_ evolution is needed to map out the molecular mechanism of C_4_ evolution. The construction of a gene regulatory network (GRN) based on the transcriptomic data constitutes an effective, reliable and powerful approach to systematically decipher the regulatory mechanism of genes either on the whole genome scale (Friedman 2004) or for a defined genes list for such a metabolic pathway (Gardner, et al. 2003; Zhang, et al. 2012). In our current study, we aimed to construct the genome-wide GRNs for four Flaveria species belonging to different photosynthetic stages (C_3_, type I C_3_-C_4_, type II C_3_-C_4_ and C_4_). The GRNs were developed based on *de novo* generated RNA-seq data derived from experimental conditions that have previously been reported to affect the C_4_ photosynthesis such as low CO_2_ (Sage 2001; Li, et al. 2014), high light (Ubierna, et al. 2013) and exogenous abscisic acid (ABA) treatment (Ueno 2001). With these GRNs, we systematically study the co-regulated TFs profiling with the C_4_ core metabolic genes (C_4_GRN) and how the C_4_GRN was re-reshuffled along the C_4_ photosynthesis evolution. This report provides a first glance on the gradual changes of GRN for the C_4_ photosynthesis-related genes in one genus characterized by its short history in C_4_ photosynthesis evolution and offers new insights about the metabolic pathways of the intermediate species.

## Results

### Transcript assembly and annotation of TFs and C_4_ genes in Flaveria species

To identify the TFs co-regulating the C_4_ genes, we constructed GRNs for the four Flaveria species based on the next-generation sequencing (NGS) of RNA-seq data. The species investigated herein represent four different photosynthetic types: *F. robusta* (C_3_; thereafter, *Fro*), *F. sonorensis* (type I C_3_-C_4_; thereafter, *Fso*), *F. ramosissima* (type II C_3_-C_4_; thereafter, *Fra*) and *F. trinervia* (C_4_; thereafter, *Ftr*) according to (Edwards and Ku 1987). Although, they belong and coexist in the same clade of the phylogenetic tree, the four species represent four different stages of the C_4_ evolution (fig. 1*A*). The RNA-seq datasets were obtained from plants subjected to treatments that were earlier reported to regulate the expression levels of the C_4_ photosynthesis-related genes such as low CO_2_ (Sage 2001; Li, et al. 2014), high light (Ubierna, et al. 2013) and exogenous ABA application (Fischer, et al. 1986; Duarte, et al. 2019). In total, between 22 and 28 RNA-seq datasets were used to construct the GRN for each species (table 1).

**Figure 1.**
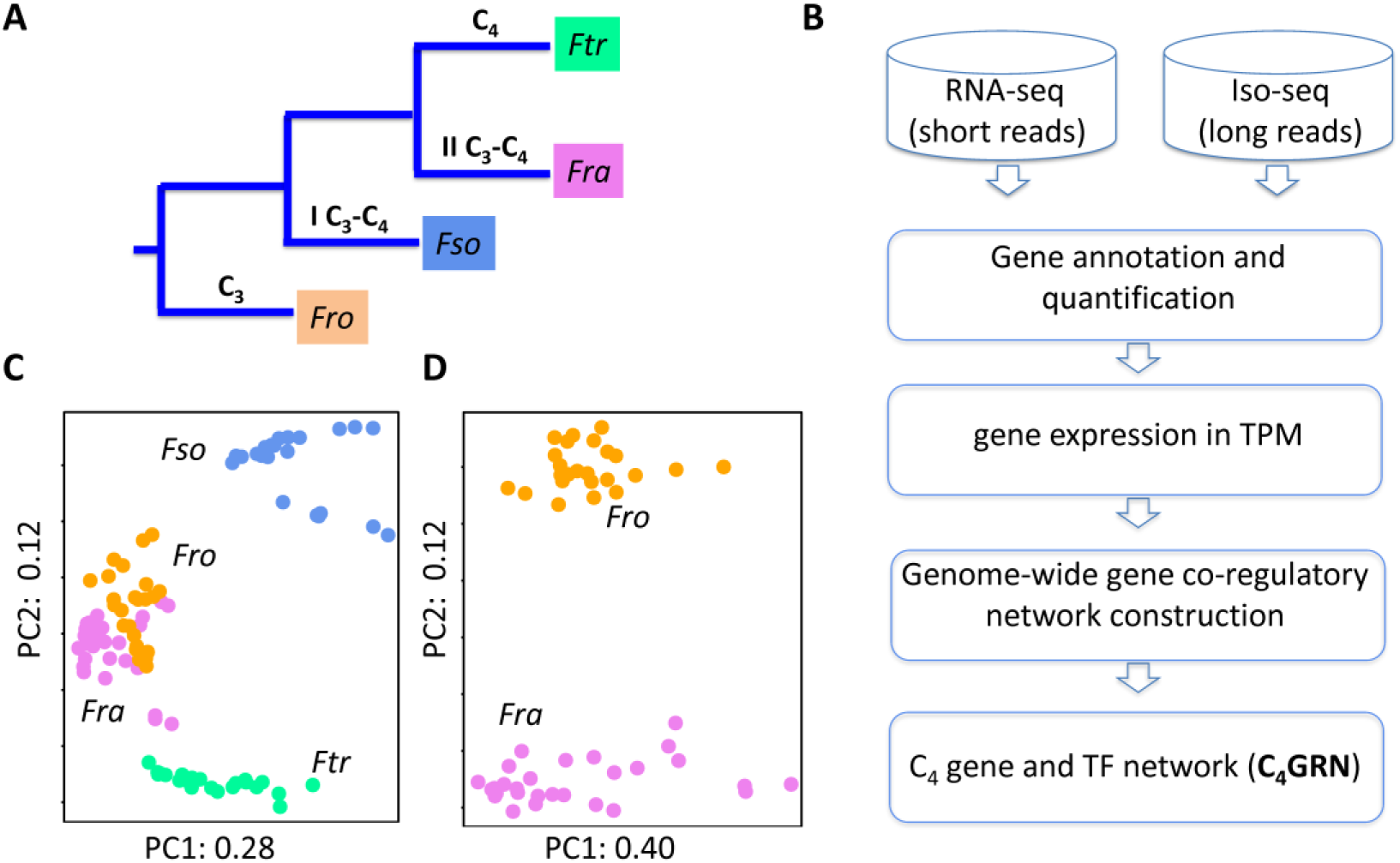
Sample information and flowchart of constructing gene co-regulatory network. (A) A schematic representation of the evolutionary relationships between four Flaveria species and their respective photosynthetic types studied herein. These are C_3_, type I C_3_-C_4_ (I C_3_-C_4_), type II C_3_-C_4_ (II C_3_-C_4_) and C_4_. (B) Flowchart of constructing gene co-regulatory network (GRN) based on RNA-seq data. In addition to RNA-seq data, we also generated a paralleled Iso-seq for *Fra* to improve gene annotation for Flaveria species. Thus, we constructed a genome-wide GRN using the conditional mutual information (CMI), and then we extracted the sub-network which contains C_4_ genes and their positively co-regulatory TFs, termed thereafter as C_4_GRN. (C-D) Principle plots of all the RNA-seq data set based on gene expression profiling. *Fra* and *Fro* clustered together when all species were plotted simultaneously (C); however, they are separated when solely their RNA-seq data were plotted (D). Abbreviations: *Fro*: *F. robusta*, *Fso*: *F. sonorensis*, *Fra: F. ramosissima*, *Ftr*: *F. trinervia*, GRN: gene co-regulatory network.

**Table1.**
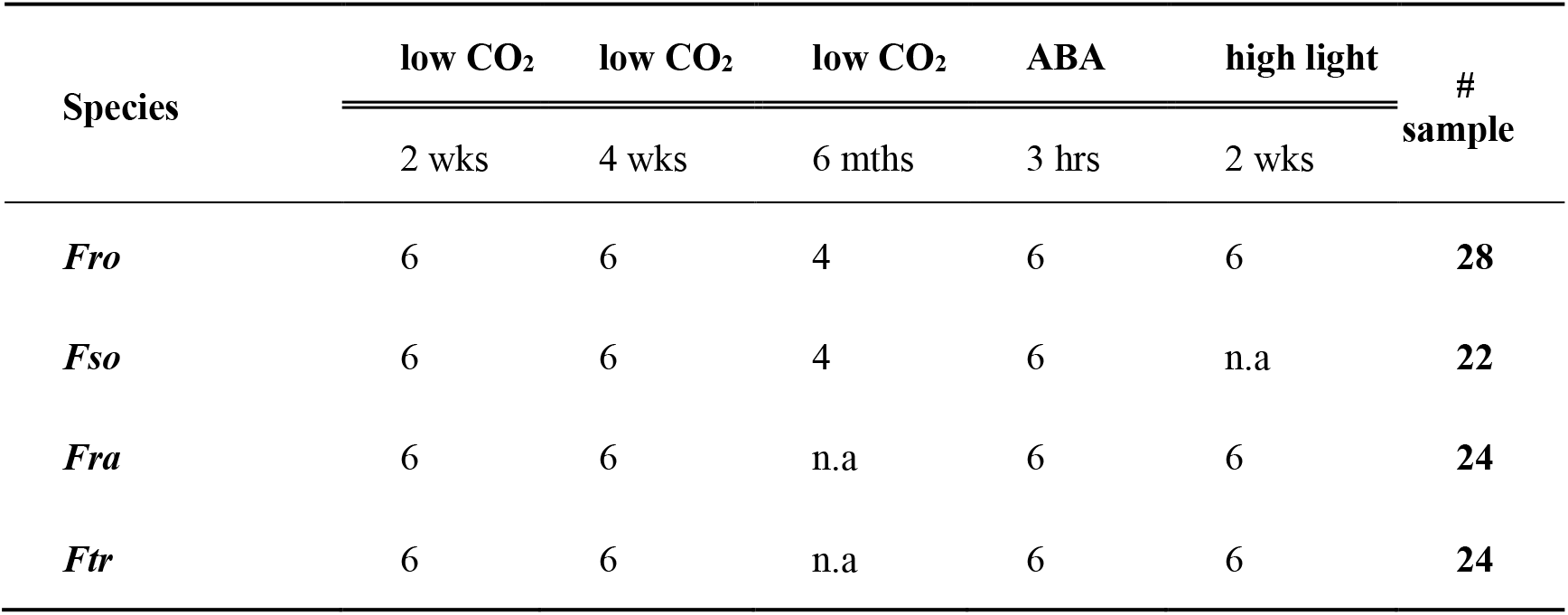
RNA-seq data generated for constructing GRN; hrs, wks and mths stand for hours, weeks and months, respectively.

Given that none of the Flaveria species has not yet a published genome sequence so far, the gene annotation for Flaveria species was mainly based on the proteins sequences in *Arabidopsis thaliana* (*A. thaliana*) published genome database (Gowik, et al. 2011; Mallmann, et al. 2014; Lyu, et al. 2015), which could not effectively distinguish the different paralogs. Hence, in order to identify the C_4_ version of C_4_ metabolic genes, we conducted full-length transcript sequences across the transcriptome of *Fra* using Single Molecular Real Time (SMRT) long read isoform sequencing (Iso-seq) of Pacific Biosciences (PacBio). The Iso-seq helps achieve full-length transcripts and effectively segregates the different paralogs. We then used the assembled gene sequences from the Iso-seq for *Fra* as a reference dataset to annotate the assembled transcripts for the three other species (fig. 1*B*). In total, 9.8 million Iso-seq long reads (LR) were obtained for *Fra*. Subsequently, we corrected the LR with next generation RNA-seq short reads (SR), which resulted in 276,323 transcripts (supplementary fig. S1A).

Thereafter, we assembled the transcripts by combining both LR and SR together. Totally, 515,464 transcripts were found; among which 99,565 were predicted to be protein-coding transcripts with an open reading frame (ORF) no less than 300 bp length. After removal of redundant sequences, 34,045 *Fra* genes were preserved and used later for further analysis in our current study. The *Fra* genes were annotated by searching their orthologs from Uniprot, which comprises 31,708,455 entries from Uniref50 and 558,898 entries from Swissprot (Supplementary Material SF2). By seeking for the orthologs in the database PlantTFDB (Jin, et al. 2017), 3,021 TFs from 57 families were annotated and characterized in *Fra* (Supplementary Material SF2). Among them, 252 TFs belong to bLHL, which represents the largest TF family in *Fra*, and 216 TFs belong to ERF (supplementary fig. S1B). In line with our findings, BLHL and ERF represent also the top two most abundant TF families in many other plants, such as *A. thaliana*, *Nicotiana tabacum*, *Oryza sativa* and *Zea mays* (*Z. mays*), as earlier well documented in the PlantTFDB website (http://planttfdb.cbi.pku.edu.cn/).

On one hand, based on the *de novo* assembled transcripts from the SR for the four species, between 66,234 and 80,154 transcripts were obtained. To confirm the accuracy of assembling, we further corroborated that around 80 to 96.8% of SR can be mapped to assembled transcripts (Supplementary Material SF3). On the other hand, 99% of the *de novo* assembled transcripts were predicted to be protein-coding ones (transcripts). Thus, around 60 to 71% of these protein-coding transcripts possess orthologs genes in *Fra* (supplementary table S1).

We then calculated the gene expression based on NGS in Transcripts per kilobase Per Million mapped reads (TPM) (Supplementary Material SF4). The principle component analysis (PCA) of the gene expression profiling revealed that *Ftr* and *Fso* are different from the remaining two other species, with the first two components exhibiting 40% of total variance. *Fro* and *Fra* could not be separated based only on their first two PCA components (fig. 1*C*). However, if only these two species were analyzed using PCA, they can be obviously distinguished into two clusters using the first two components, which reflect a total variance of 52% (fig. 1*D*).

Since most of the C_4_ genes belong to multiple-gene families (Moreno-Villena, et al. 2018), it is interesting to determine the C_4_ version of C_4_ genes. Hence, we manually selected the C_4_ version of C_4_ genes based on two criteria. (1) The C_4_ version gene should show higher transcript abundance compared to other paralogs; (2) The C_4_ version should exhibit higher transcript abundance in C_4_ species than C_3_ ones. Predominately, the C_4_ version could mainly be selected based on the first criterion. For instance, two paralogs of *β*-carbonic anhydrase 1 (*β*CA1) were annotated in the C_4_ species *Ftr* with a single paralog (Gene00137) having a transcript abundance of around 20,000 TPM and another paralog (Gene23895) having a transcript abundance of around 1,000 TPM. Besides, the Gene00137 showed higher transcript abundance in the C_4_ species *Ftr* than in the C_3_ species *Fro* (supplementary fig. S2A-D). Accordingly, the Gene00137 was selected to be the C_4_ version of CA1 in *Ftr*. The latter (*Ftr*) uses a typical NADP-dependent malic enzyme (NADP-ME) type C_4_ photosynthesis, where only NADP-ME was up-regulated in *Ftr* compared to *Fro*, while NAD-dependent malic enzyme (NAD-ME) and PEP carboxykinase (PEPCK) showed comparable transcript abundance in these four species (supplementary fig. S2A-D). Except the enzymes and transporters required for the NADP-ME pathway, we found as well a significantly higher transcript abundance of alanine aminotransferase (AlaAT) in *Ftr* if compared to *Fro*; however, aspartate aminotransferase (AspAT) did not show such a pattern between the four species. We, therefore, included this gene (AlaAT) in the list of C_4_ core metabolic genes. The AlaAT is actually known to transfer ammonia from BSC to the MC by glutamate/2-oxoglutarate shuttle in order to maintain the nitrogen homeostasis between the two compartments, MC and BSC (Mallmann, et al. 2014). All the C_4_ core metabolic genes show higher transcript abundance in the C_4_ species *Ftr* than that in the C_3_ species *Fro* and such patterns are found to be conserved under the four experimental conditions used in this investigation (supplementary fig. S2). Among the different treatments, four weeks low CO_2_ treatment shows the substantially largest effect on the C_4_ genes expressions for the four species (supplementary fig. S3).

To confirm the accuracy of the transcript quantification via RNA-seq data processing, a qRT-PCR analysis was performed for 14 genes on the same RNA samples used for RNA-seq in plants exposed for four weeks to low CO_2_ of 100 ppm (supplementary fig. S4). Thus, we found an averaged Pearson correlation coefficient of 0.78 between transcript abundance from RNA-seq and that (transcript abundance) from qRT-PCR analysis. This shows the reliability and accuracy of our transcript quantification via RNA-seq data processing. The expression patterns of C_4_ core metabolic genes obtained from qRT-PCR are consistent with those evaluated using RNA-seq data (fig. *2A* and *B*). Indeed, figure 2*B* depicts the proposed C_4_ core metabolic pathway based on the identified C_4_ genes in Flaveria, which including 8 enzymes: CA1, PEPC1, PEPC-K, NADP-dependent malate dehydrogenase (NADP-MDH), NADP-ME4, AlaAT2, pyruvate/orthophosphate dikinase (PPDK) and PPDK regulatory protein 2 (PPDK-RP2), and 5 transporters: dicarboxylate transporter 1 (DiT1), dicarboxylate transporter 2 (DiT2), bile acid sodium symporter 2 (BASS2), sodium: hydrogen antiporter 1 (NHD1) and phosphate/phospho*enol*pyruvate translocator 1 (PPT1).

**Figure 2.**
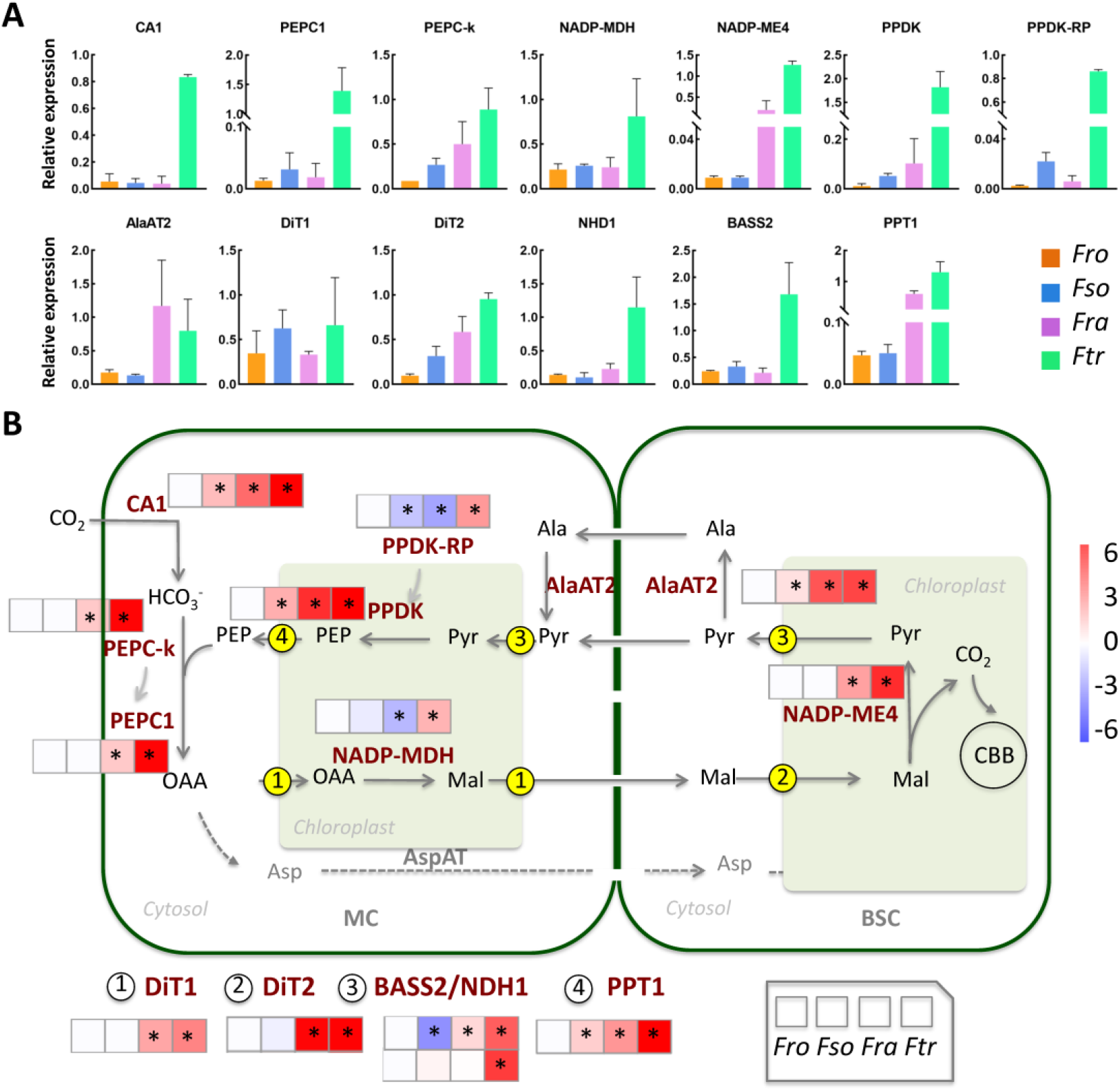
C_4_ core metabolic pathway and its involved genes. (A) The histogram plots depict the quantification of the determined C_4_ version of C_4_ genes based on qRT-PCR analysis performed on leaf samples of plants grown at low CO_2_ concentration (100 ppm) for four weeks. (B) The diagram of the core C_4_ pathway and the involved genes in Flaveria. *Ftr* is a typical NADP-ME type of C_4_ photosynthesis. The heatmaps show the log2-transformed fold changes of transcript levels of C_4_ genes assessed as Transcript per kilobase Per Million mapped reads (TPM) in each species compared with their counterparts in the C_3_ species *Fro*. RNA-seq quantification from 4-week low CO_2_ experiment is showed here. A star (*) represents significant changes at a *P*-value < 0.05, and FC > 1.5. Abbreviations: CA1, carbonic anhydrase 1; PEPC1, phospho*enol*pyruvate carboxylase 1; PEPC-K, PEPC kinase; NADP-MDH, NADP-dependent malate dehydrogenase; NADP-ME4, NADP-dependent malic enzyme 4; AlaAT2, alanine aminotransferase 2; PPDK, pyruvate/orthophosphate dikinase; PPDK-RP: PPDK regulatory protein. Metabolite transporters were displayed in the encircled numbers located on the membrane, with the consecutive number from 1 to 4 refer to DiT1 (dicarboxylate transporter 1), DiT2 (dicarboxylate transporter 2), BASS2 (bile acid sodium symporter 2), NHD1 (sodium: hydrogen antiporter 1) and PPT1 (phosphate/phospho*enol*pyruvate translocator 1), respectively. Species abbreviations are as depicted in Fig. 1.

### The properties of C_4_GRNs in four Flaveria species

For this purpose, we constructed a genome-wide GRN based on the RNA-seq data for each species. We further built a “C_4_GRN” which retains only the C_4_ genes and their positively co-regulated TFs. We termed this proposed network as “C_4_GRN” and designated the TFs involved in C_4_GRN as “C_4_TFs” (Supplementary Material SF5). The genome-wide GRN of type I C_3_-C_4_ species *Fso* contains the minimum genes (14,677 genes) and the maximum interactions (7,841,065 interactions). In contrast, the C_4_ species *Ftr* contains 14,761 genes and 5,769,379 interactions (fig. 3*A*). Similarly, the C_4_GRN of *Fso* includes 220 TFs, followed by the networks of *Ftr* and *Fro* and *Fra* with 196, 185 and 114 TFs, respectively (fig. *3A*). In the same regard, more than half of the identified C_4_TFs are found to be specific for each species. For instance, 57.9% C_4_TFs of *Fra* is not shared with any other species and the percentage of species-specific C_4_TFs in *Fro* is 69.2% (fig. *3B*).

**Figure 3.**
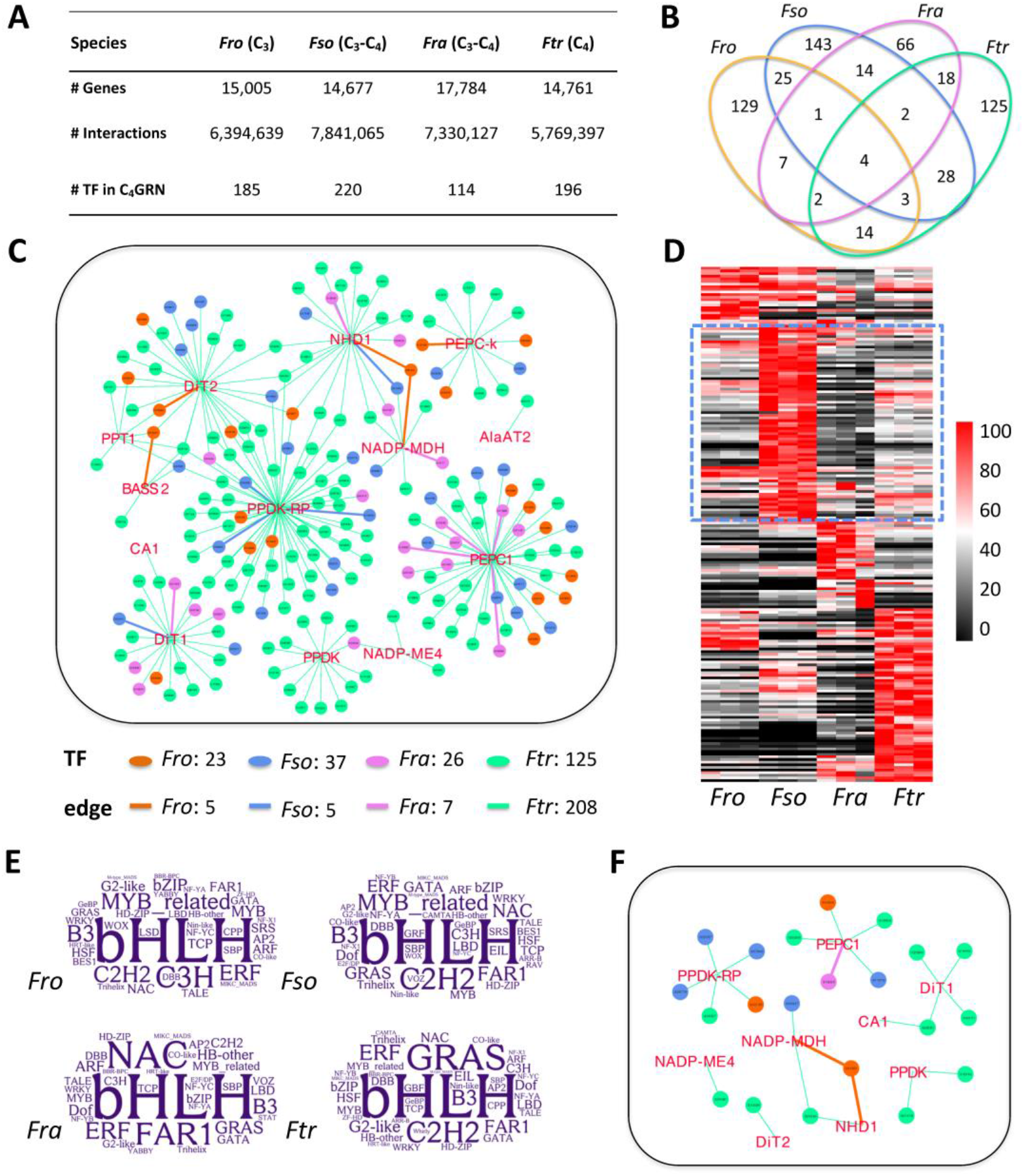
The C_4_GRN properties. (A) Basic information on the genome-wide GRN and C_4_GRN in four Flaveria species. (B) The overlapped TFs in the C_4_GRN between the four investigated Flaveria species. (C) The C_4_GRN in the C_4_ species *Ftr* was displayed, where only the interactions between the C_4_ genes and their co-expressed TFs are shown. The C_4_ genes were depicted in red font and the TFs were shown in circles with 4 different colors. The overlapped TFs with other three species were exhibited in different colors as depicted by the legend “TF” below the panel. The overlapped C_4_ genes-TFs interactions were displayed in the same colors used for the overlapped TFs as reflected by “edge” in the legend. (D) The expression patterns of the 196 co-regulated TFs with the C_4_ genes in *Ftr* were exhibited in a heatmap, which was calculated based on the RNA-seq data obtained from plants, of each species, experienced low CO_2_ (100 ppm) for four weeks. Colors represent the relative transcript abundance, with red, white and black mean high, median and low expression level, respectively. The comparison of the TFs with the highest transcript abundance in *Fso* to those of the three other species was delimited by the selected zone with the blue dashed-edge rectangle on the heatmap. (E) Word clouds showing the C_4_TFs frequency in different TF families, with large word size indicates the higher frequency. (F) Co-regulation network of C_4_ genes and TFs from GRAS family in *Ftr*. The colors of circles and edges are same as in the legend of panel (C). Species abbreviations are the same as Fig. 1.

In total, 36.2% C_4_TFs in *Ftr* overlapped with those of at least one of the three other species. The C_4_GRN of *Fso* exhibits 37 TFs overlapped with *Ftr*, displaying the highest overlap degree with *C*_4_ species compared to *Fro* and *Fra* (fig. *3C*). However, the co-regulated C_4_ genes of these overlapped C_4_TFs changed in these different Flaveria species. Five C_4_ genes-TFs co-regulations are common between *Fro* and *Ftr*, involving five C_4_ genes, which are NHD1, NADP-MDH, DiT2, BASS2 and PEPC-k. The same number of C_4_ genes-TFs co-regulations exists between *Fso* and *Ftr*, among them three involving the three following genes: PPDK-RP, DiT1 and NHD1. Notably, *Fra* shares seven C_4_ genes-TFs co-regulations with *Ftr* with four engaging PEPC1 (fig. *3C*). The other three co-regulations targeted three other genes including DiT1, NHD1 and NADP-MDH.

In the C_4_GRN of *Ftr*, PPDK-RP shows the greatest number of co-regulated TFs with 63 TFs, followed by PEPC1 with 45 TFs, while CA1 ranks the latest with only one TF (fig. *3C*). Interestingly, 74 C_4_TFs (37.8%) in *Ftr* show the highest transcript abundance in *Fso* among the four species, even more than the number of C_4_TFs showing the highest transcript abundance in *Ftr* (62 TFs), which represents 31.6% from the total C_4_TFs of *Ftr* (fig. 3*D*).

The most abundant C_4_TFs in the four species is bHLH (fig. *3E*). The next most abundant TFs in both *Fro* and *Fso* is MYB-related TFs, whereas *Fra* has NAC family TFs being the second most prevailing TFs; however, GRAS family TFs is the second most abundant, after bHLH, in the *Ftr*. In fact, 21 C_4_TFs belonging to the GRAS family in *Ftr* are co-regulated with 9 over 13 C_4_ genes (fig. *3F*).

Therefore, compared to C_3_ and both types of the intermediate species, the C_4_ species *Ftr* recruited a large number of TFs, which are co-regulated with PEPC1 or PPDK-RP. Simultaneously, several TFs from the GRAS family were also recruited. *Fso* shares the largest number of C_4_TFs with the C_4_ species, while *Fra* shares the maximum C_4_ genes-TFs co-regulations with the C_4_ species, where four of them (co-regulations) are associated with PEPC1.

### Assigned functions of C_4_TFs and C_4_ genes

To gain insights on the functions of C_4_TFs in each species reported herein, we firstly investigated the enriched functions of C_4_TFs. The obtained results show that C_4_TFs from the four species consistently are enriched in glycolysis (SI6). The C_4_TFs from *Fso* and *Fra* show more enriched GO functions than *Fro* and *Ftr*. In *Fso*, C_4_TFs are also enriched in histone acetyltransferase activity and brassinosteroid mediated signaling pathway (Supplementary Material SF6); whereas, in *Fra*, C_4_TFs are enriched in functions related to glycine mentalism, *i.e*., glycine dehydrogenase (decarboxylating) activity and glycine catabolic process. However, None C_4_TFs from all the species exhibit over-represented functions directly related to photosynthesis.

Afterwards, we investigated the related functions of C_4_TFs in each species by examining the functions of their co-regulated genes. To achieve that, we firstly clustered the whole genome-wide GRN into multiple sub-networks (modules) and then studied the biological function of the modules enriched in C_4_TFs (MET; fig. 4*A*). Specifically, 222, 363, 642 and 295 modules were identified in *Fro*, *Fso*, *Fra* and *Ftr*, respectively (fig. 4*B*). We found only a single MET in *Fro*, *Fso* and *Ftr* individually, while two METs were characterized in *Fra* (*Fisher*’s test, *P* < 0.001, BH adjusted); however, uniquely one of the METs in *Fra* showed over-represented GO functions (fig. *4B*).

**Figure 4.**
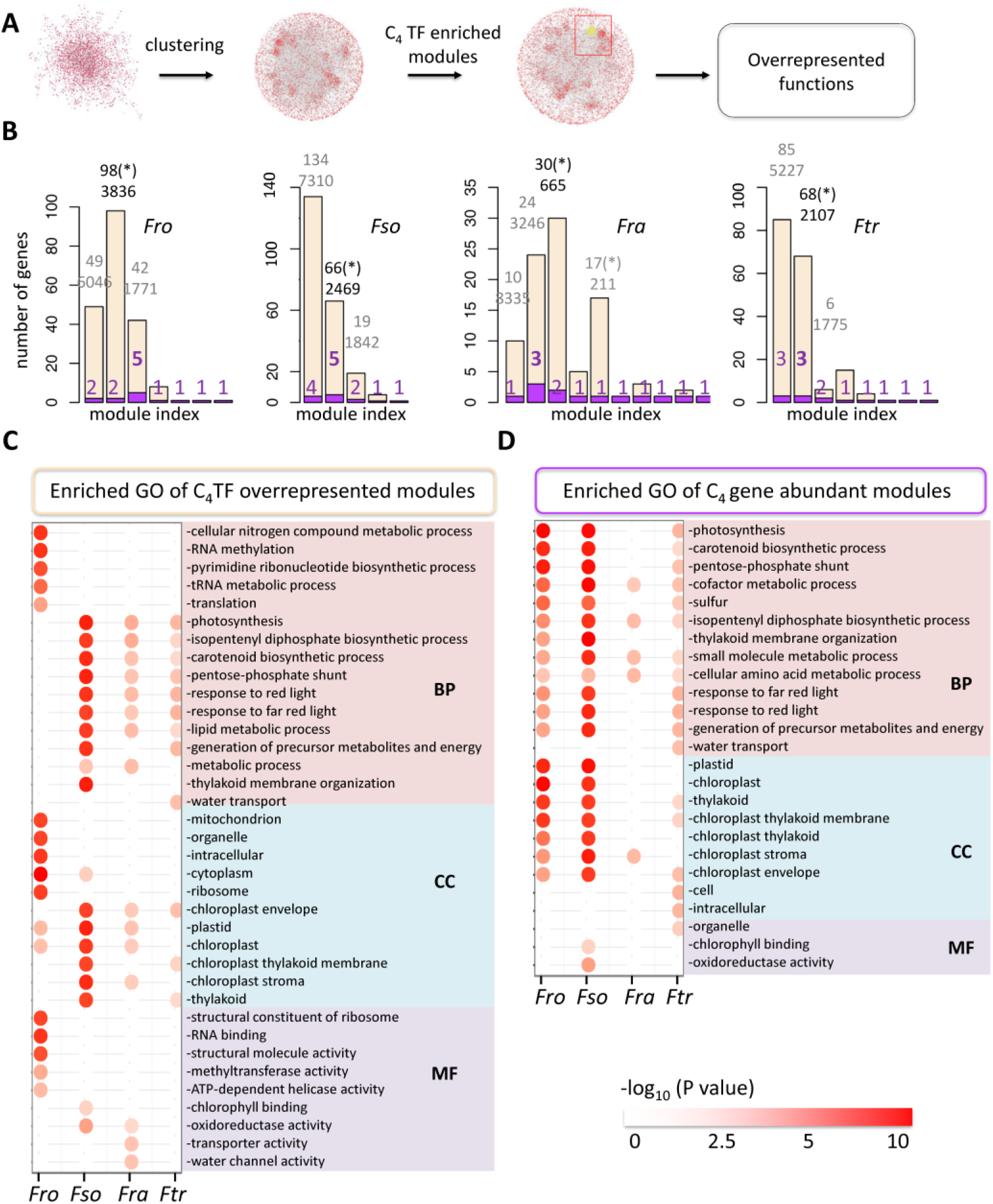
Biological functions of genes from MET and MMC_4_ in different species. (A) A predictive diagram displaying the analysis pipeline used to identify the potential functions of C_4_TFs. A genome-wide GRN was firstly clustered into modules. The latters (modules) enriched in the C_4_TFs (*Fisher’s* test, *P* < 0.001, BH adjusted) were then determined and termed modules enriched in C_4_TFs (MET). Over-represented GO functions of genes in the MET were calculated using *Fisher*’s test (*P* < 0.001, BH adjusted). (B) Bar plots show the number of C_4_TFs and C_4_ genes in the modules that encompass C_4_ genes. Bars in antique white represent the number of C_4_ TFs and bars in orchid represent the number of C_4_ genes. The number given in grey and black show the number of C_4_TFs (upper value) and total number of genes (bottom value) in each module, and the numbers in orchid show the number of C_4_ genes. The star “*” indicates that the module was a MET. Over-represented GO terms of genes from MET (C) and genes from modules encompassing the maximum C_4_ genes (MMC4) (D) were shown. Abbreviations: BP, molecular process; CC, cell component; MF, molecular function. The species abbreviations are as depicted in Fig. 1. BH means Benjamini and Hochberg.

We further examined the top five enriched GO terms of genes in METs from three aspects, *i.e*., biological process (BP), cell component (CC), and molecular function (MF). Thus, *Fro* exhibits a large proportion of GO terms from BP related to the transcriptional and post transcriptional processes, *e.g*., RNA methylation, tRNA metabolic process and translation. This species (*Fro*) includes also a single GO term associated with the cellular nitrogen compound metabolic process (fig. *4C*). In terms of CC, the enriched GO functions are mainly related to the various cellular compartments, such as mitochondrion, organelle, intracellular, cytoplasm and ribosome (fig. *4C*), but not the chloroplast.

Indeed, a substantial overlap exists, regarding the enriched GO terms in METs, between *Fso*, *Fra* and *Ftr*. Hence, many of these overlapped GO terms of BP are involved in photosynthesis, or terms directly related to photosynthesis, such as photosynthesis and thylakoid membrane organization; and some other items indirectly linked to photosynthesis, including carotenoid biosynthetic process, chloroplast relocation, response to red light and to far-red light (fig. *4C*). Two other biological processes, *i.e*., the pentose phosphate shunt and lipid metabolic pathway, are also found among the overlapped GO terms in *Fso, Fra* and *Ftr*. Consistently, almost the whole enriched GO functions in the terms of CC are predominately attributed to the chloroplast term such as chloroplast, chloroplast envelope, chloroplast thylakoid membranes, chloroplast stroma, and thylakoid (fig. *4C*).

We thereafter investigated the functions of the C_4_ orthologous genes in different species by looking after the over-represented GO terms in modules that include the maximum C_4_ orthologous gene (MMC_4_). In *Fso*, the MET and MMC_4_ are the same module, which includes five C_4_ genes, which are CA1, DiT2, NADP-ME4, NADP-MDH and PPDK. Two MMC_4_ were found in *Ftr* with one coincides with the MET in *Ftr*. A considerable fraction of the over-represented GO terms of MMC_4_ are shared among *Fro*, *Fso* and *Ftr*, and many of them are related to the photosynthesis and chloroplast (fig. *4D*). The shared GO terms, associated with BP, between *Fro, Fso*, and *Ftr*, include photosynthesis, carotenoid biosynthetic processes, pentose-phosphate shunt, cofactor metabolic process, sulfur metabolism, isopentenyl diphosphate biosynthetic processes, small molecule metabolic pathways, responses to red and far-red light, generation of precursor metabolite and energy. Furthermore, the shared GO terms, related to CC, between the three species (*Fro*, *Fso*, *Ftr*) englobe thylakoid, chloroplast membrane and envelope. Most of these enriched GO terms in *Fro*, *Fso* and *Ftr* were not shared with *Fra* (fig. *4D*).

Taken together, the orthologs of C_4_ key metabolic genes in the genus Flaveria were already co-regulated with the photosynthetic genes in C_3_ species, whereas, the C_4_ photosynthesis related regulators were widely recruited during the latest C_4_ evolution stages.

### The overlap of C_4_GRNs between the intermediate and C_4_ species

Our above analysis shows that many C_4_TFs are co-expressed with the photosynthesis related genes in two intermediate C_3_-C_4_ species (fig. 4*C*). Given the metabolic and anatomical adjustments between the intermediate species and the C_4_ ones (species), what are the missing regulatory factors related to C_4_ photosynthesis in these intermediate C_3_-C_4_ species compared to C_4_ species? On one hand, we found that the 37 C_4_TFs shared between *Fso* and *Ftr* are co-regulated with 12 of the 13 C_4_ genes (except the PEPC1, fig. 5*A*). This is consistent with the lower transcript abundance of PEPC1 in *Fso* (fig. 2*B* and supplementary fig. S2A-D). On the other hand, in the type II C_3_-C_4_ species *Fra*, 26 C_4_TFs were found to be common between *Fra* and *Ftr*, which are co-expressed with 8 over 13 C_4_ genes in *Fra* (fig. 5*B*). Four among the 26 TFs are co-regulated with PEPC1 (fig. 3*C* and Supplementary Material SF 5).

**Figure 5.**
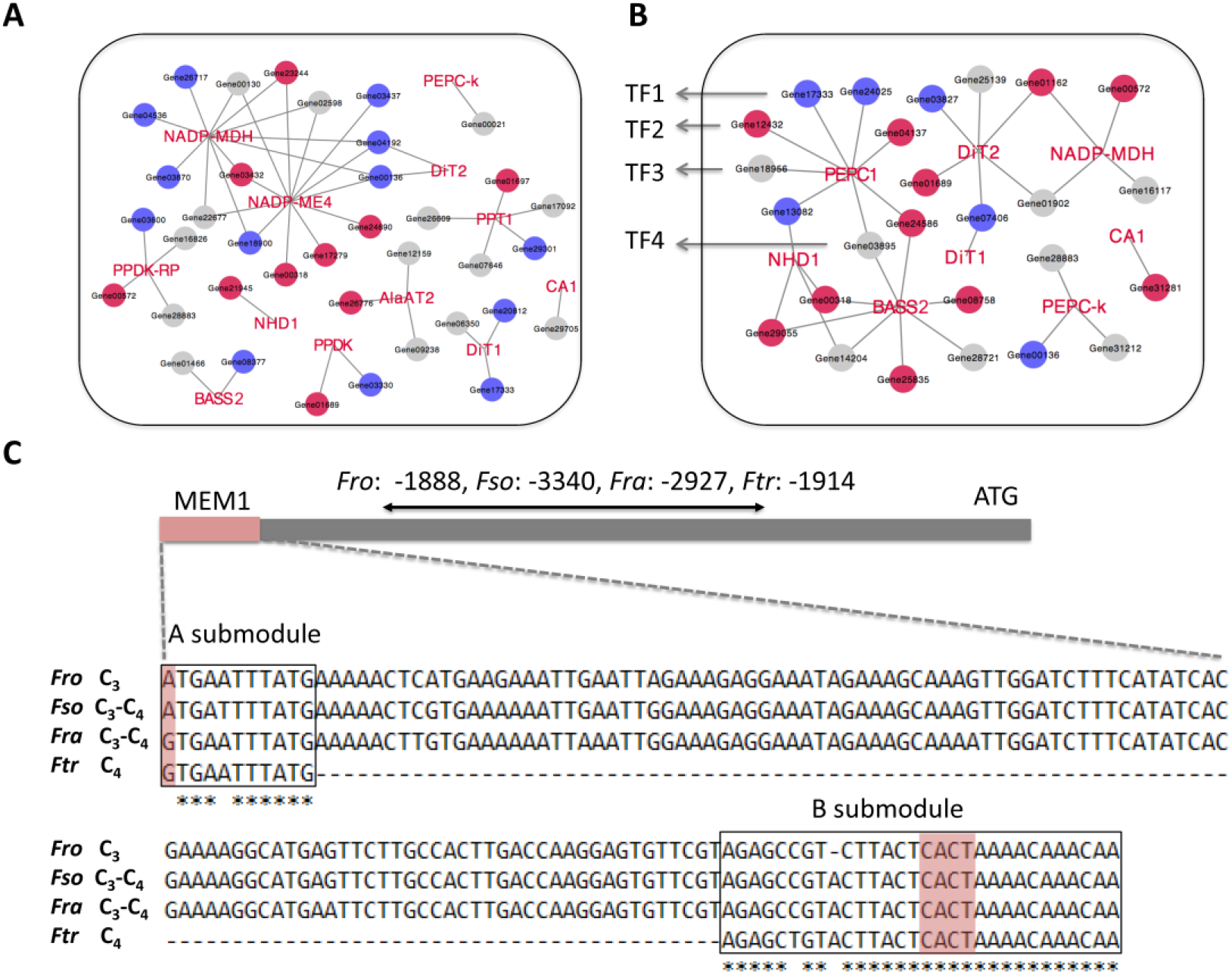
The C_4_GRN overlap between the intermediate and C_4_ species. (A) and (B) show the sub-networks of C_4_GRN in *Fso* and *Fra*, respectively. These show the regulatory relationship between C_4_ gene and the shared C_4_TFs with *Ftr*. The C_4_ genes are depicted in red font. TFs are shown in circles, with red, blue and grey colors represent the TFs showing an increased, decreased, or similar (comparable) transcript abundance in the C_4_ species *Ftr* compared to that in the intermediate species (edgeR, P < 0.05 and FC > 1.5). Four TFs co-regulating the PEPC1 are found to be conserved between *Fra* and *Ftr*, as labeled by TF1-4, which is from TF family of MYB_related, NF-YA, GRAS and DBB, respectively. (C) The mesophyll expression module 1 (MEM1) sequences of the PEPC1 promoters from four species. The A and B submodules are highlighted in boxes. Asterisks show identical nucleotides in the two modules. Red zones indicate the single nucleotide difference in the A submodule, and the required tetranucleotide CACT in the B submodule. The full promoter sequences are available in Supplementary Material SF7. Abbreviations of enzymes are same as those given in Fig. 2.

A mesophyll expression module 1 (MEM1) located on the PEPC1 promoter region of the C_4_ species *F. bidentis* and *Ftr* was reported to control the mesophyll specific high expression of PEPC1 (Gowik, et al. 2004). We examined, herein, whether the MEM1 is present in the PEPC1 promoter region of *Fra*. Indeed, the MEM1 is C_4_ type in *Fra*, while it is a C_3_ type in *Fro* and *Fso* (fig. 5*C* and Supplementary Material SF7), which is in consistency with the up-regulation of PEPC1 expression in *Fra* compared to *Fro* and *Fso* (fig. 2*B* and supplementary fig. S2). The four TFs mentioned above to be co-regulated with PEPC1 in both *Fra* and *Ftr* (fig. 5*B*) are involved or annotated as a negative regulation function (Gene03809), a TF belongs to NF-YA7 family (Gene12432), regulation of drought-responsive genes (Gene17333) and repressor of gibberellins response (Gene18956, supplementary table S2).

The shared 37 TFs between *Fso* and *Ftr* are mainly enriched in two GO functions, proton-exporting ATPase activity (phosphorylative mechanism) and other cellular processes (Supplementary Material SF6). However, the shared 26 TFs between *Fra* and *Ftr* are enriched in GO functions of glycolysis and chloroplast membrane (Supplementary Material SF6). Compared to *Fso*, *Ftr* newly gained 159 TFs and lost 183 TFs (fig. 3*B*). The newly recruited TFs are enriched in GO functions of glycolysis, chloroplast, nucleus and coenzyme A metabolic process. Otherwise, the lost TFs (present in *Fso* but not in *Ftr*) are enriched in glycolysis, mitochondrion, histone acetyl-transferase activity and isocitrate dehydrogenase (NAD^+^) activity (Supplementary Material SF6). Compared to *Fra*, *Ftr* newly grained 170 TFs and lost 88 TFs. The gained TFs are enriched in GO functions of nucleus, coenzyme A metabolic process and hydroxyl-methyl-glutaryl-CoA reductase (NADPH) activity; whereas, the lost TFs (present in *Fra* but not in *Ftr*) are essentially enriched in GO items of glycine dehydrogenase (decarboxylating) activity and glycine catabolic process (Supplementary Material SF6). Therefore, C_4_TFs in the two intermediate species as well as C_4_ species are involved in divergent functions.

Taken together, it seems that the mechanisms underlying the mesophyll specific high expression of PEPC1 are not yet recruited in *Fso*. However, in *Fra*, the mechanisms governing the cell-specific high expression of PEPC1 appears, at least, partially recruited. Nevertheless, five C_4_ genes including PPDK, NADP-ME4, PPDK-RP, AlaAT2, PPT1 are exempted from the co-regulation process by the shared C_4_TFs between *Fra* and *Ftr*, which is in agreement with the difference between the enriched GO terms in MMC_4_ for *Fra* compared to those for *Ftr* (fig. *4D*).

### The evolution of C_4_GRN towards a C_4_ photosynthesis stage

In this section, we illustrated how the C_4_GRN of C_4_ species gradually evolved during the evolution process. We firstly defined the C_4_ required TFs (thereafter, C_4_ReTFs) as C_4_TFs in *Ftr* that show significantly higher transcript abundance in *Ftr* than in the C_3_ species *Fro* (*P* < 0.05 and fold change (FC) > 1.5). Eventually, we identified 93 C_4_ReTFs, targeting all the 13 C_4_ genes in *Ftr* (fig. 6*A*).

**Figure 6.**
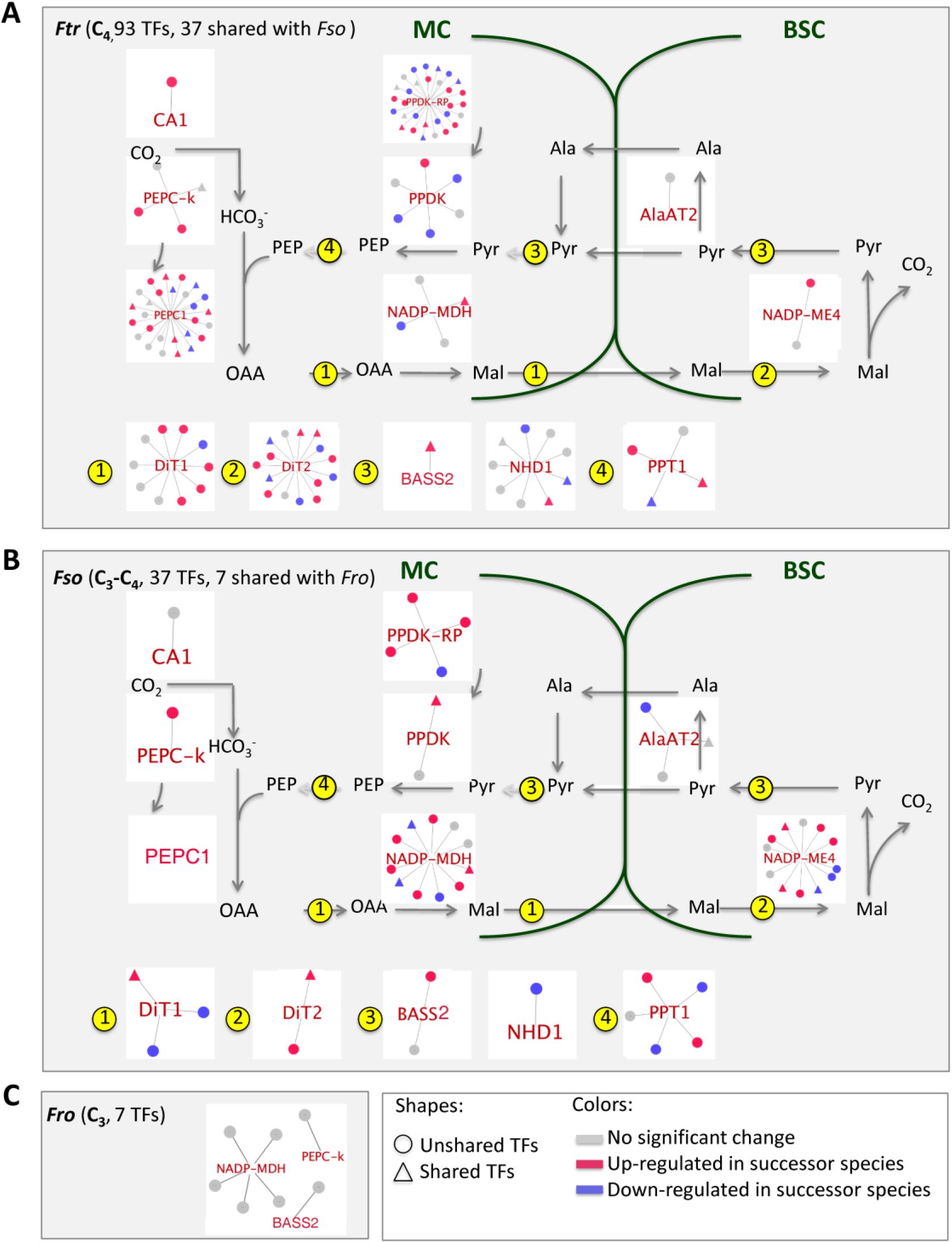
The C_4_GRN evolution towards a C_4_ photosynthesis stage. (A) In *Ftr*, 93 C_4_GRN TFs show higher transcript abundance than *Frob* (edgeR, P < 0.05 and FC > 0.5), termed as C_4_-required TFs (C_4_ReTFs), which co-regulate all the 13 C_4_ core metabolic genes. (B) 37 of C_4_ReTFs are present in the C_4_GRN of *Fso*, co-regulated with all C_4_ core metabolic genes except PEPC1. TFs are shown in circles and triangles, with triangles presenting TFs shared with either *Fso* (panel A) or *Fro* (panel B). Red, blue, and grey colors represent TF showing higher, lower and similar transcript abundance compared to *Fso* (A) or *Fro* (B) (edgeR, P < 0.05 and FC > 1.5). (C) Seven among the 37 TFs present in the C_4_GRN of *Fro* which (seven TFs) co-regulate three orthologs of C_4_genes. The C_4_GRN of *Ftr* and *Fso* are shown in Fig. S5. Abbreviations of species and enzymes are same as those given in Fig. 2.

In *Fso*, all the shared 37 C_4_TFs with *Ftr* are included in the 93 C_4_ReTFs (fig. *6B*). We classified the modified C_4_ReTFs between *Fso* and *Ftr* into two types (fig. 7*A*). The first one includes the existing C_4_ReTFs in *Fso*, which exhibited enhanced expressions in *Ftr*. We found that 10 out of the 37 C_4_ReTFs (fig. 6*A* and *B*) displayed higher transcript abundance in *Ftr* than in *Fso* (P < 0.05 and FC > 1.5), with five of them being co-regulated with PEPC1 in *Ftr* (fig. 6*A*). The second one (type) encompasses the absent C_4_ReTFs from the C_4_GRN of *Fso*. In this latter type, we identified 56 C_4_ReTFs (fig. 6*A*). Besides, new co-regulations were developed between the first type of C_4_ReTFs and the C_4_ genes during the transition from type I C_3_-C_4_ to C_4_ species (fig. 7 and supplementary fig. S5).

**Figure 7.**
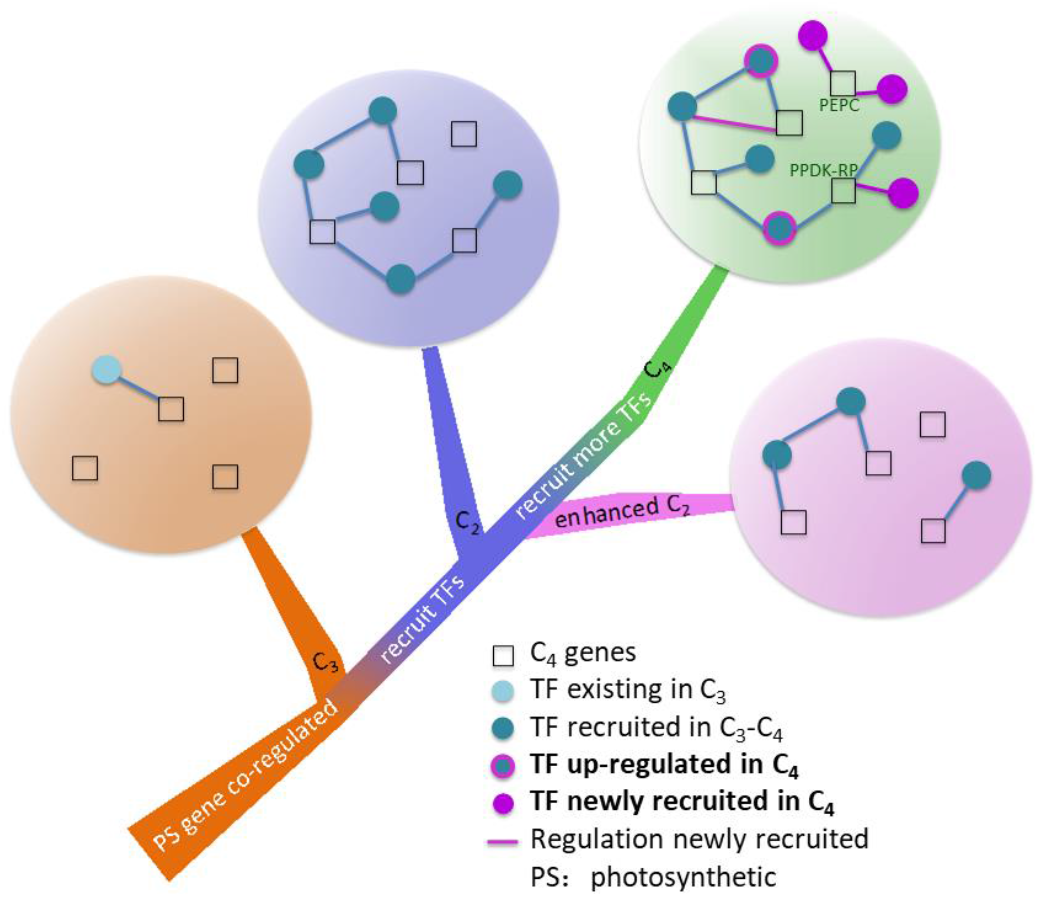
A proposed model for the C_4_GRN evolution in *Flaveria* genus. Two key stages are involved during the C_4_GRN evolution in Flaveria genus. Though the C_4_ genes orthologs were co-regulated with the photosynthesis related genes in C_3_ species, the TFs required for the C_4_ photosynthesis remain still absent. During the transition from the C_3_ species to type I C_3_-C_4_ species, a large number of C_4_-required TFs (C_4_ReTFs) were recruited, but not for all C_4_ genes. The second stage constitutes the transition from type I C_3_-C_4_ species to C_4_ species, where a complete C_4_GRN establishment could be achieved. This C_4_GRN includes the recruitment of new TFs, enhancement of the transcript abundance of C_4_ReTFs that are already recruited in type I C_3_-C_4_ species and the formation of new regulations between the already pre-existing TFs and other C_4_ genes. *Fra* may represent a divergent CO_2_ CCM as C_4_ species, in which the C_4_GRN is different from those of type I C_3_-C_4_ and C_4_ species. PEPC and PPDK-RP show a greater number of co-regulated TFs than other C_4_ core metabolic genes in *Ftr*.

Seven of the 37 C_4_ReTFs recruited in *Fso* are present in the C_4_GRN of *Fro* and co-regulate with three C_4_ core metabolic genes in *Fro*, *i.e*., NADP-MDH, BASS2, PEPC-k (fig. 6*C*). Six over the seven TFs exhibit significantly higher transcript abundance in *Fso* than in *Fro* (P < 0.05 and FC > 1.5; fig. 6*B*). Among the 30 C_4_ReTFs that were newly recruited in *Fso*, 17 are up-regulated in *Fso* if compared to C_3_ species (P < 0.05 and FC > 1.5; fig. 6*B*).

## Discussion

A comparative analysis of the GRN between species could shed light on the driving force and the mechanisms underlying the adaptation and evolution processes (Thompson, et al. 2015). In this study, we provide a comprehensive survey about the changes of GRN in four species from the genus Flaveria, representing four different stages in C_4_ evolution, *i.e*., *Fro* (C_3_), *Fso* (type I C_3_-C_4_), *Fra* (type II C_3_-C_4_) and *Ftr* (C_4_) (Edwards and Ku 1987). Considering that the transition from C_3_ to C_4_ photosynthesis spans a short evolutionarily divergence time (Christin, et al. 2011), the comparison between the GRNs for these four species could shed new light on the evolution of C_4_ photosynthesis.

### Fso recruited a large number of C_4_ReTFs very likely through reorganizing pre-existing cis-elements

*Fso* (type I C_3_-C_4_) shares the highest number of C_4_TFs overlapping with *Ftr* (C_4_), with a number of 37 (fig. 3*B*), which were predicted to regulate 12 of the 13 C_4_ core metabolic genes studied in this work (fig. 5). All the 37 TFs were characterized as C_4_ReTFs since they show higher expression levels in the C_4_ *Ftr* than in the C_3_ *Fro* (fig. 6*B*). Besides, the MET and MMC4 coincide to form the same module in *Fso*, which exhibits a large fraction of the enriched GO functions related to photosynthesis (fig. 4). This is in agreement with the current notion that the metabolism of C_3_-C_4_ species, which performs C_2_ CCM, might represent a metabolic pre-condition for the emergence of C_4_ photosynthesis, since many of the involved enzymes and transporters in this poise are common with those in C_4_ photosynthesis (Mallmann, et al. 2014). Four of the 37 TFs were annotated to be stress response related (Gene02598 and Gene29705: DBB family, salt stress; Gene04536: ERF family, regulation of expression by stress factors and Gene29301: bHLH family, oxidative stress, Supplementary Material SF8). This is in line with earlier reports that C_2_ CCM evolved to cope with conditions favoring photorespiration (Sage, et al. 2012; Lundgren and Christin 2017). The large number of the recruited C_4_ReTFs in *Fso* might be accomplished by recruiting pre-existing *cis*-elements and/or rewiring pre-existing TFs to gain new functions (Brown, et al. 2011; Reyna-Llorens, et al. 2018).

In this analysis, we found that 30 over the recruited 37 C_4_ReTFs are not represented in the C_4_GRN of *Fro*, C_3_ (fig. 6*C*). These TFs might be recruited through hiring the already existing *cis*-elements into C_4_ genes. In this regard, it has been reported that the C_4_ genes from *Z. mays*, C_4_, harbor a number of conserved *cis*-elements with their counterparts in *Oryza sativa*, C_3_ (Xu, et al. 2016). Furthermore, multiple C_4_ genes may recruit the same (or similar) *cis*-elements. For example, PEPC1 and CA3 (named CA1 in this study) harbor MEM1 and MEM1-like motif that determine the MC expression specificity in the C_4_ species *F. bidentis* (Akyildiz, et al. 2007; Gowik, et al. 2017); CA and PPDK share a MEM2 motif in 5’UTR in the C_4_ species *Gynandropsis gynandra* (*G. gynandra*) which controls the expression specificity in the MC (Williams, et al. 2016). In this context, it has been also reported that *cis*-elements from the C_4_ genes could be recognized by the TFs of C_3_ species (Ku, et al. 1999; Nomura, et al. 2000; Fukayama, et al. 2001; Akyildiz, et al. 2007). In the case of NAD-ME, the *cis*-element controlling the BS specific expression in the C_4_ species *G. gynandra* similarly shows the same cell specificity in the C_3_ species *A. thaliana* (Brown, et al. 2011). Furthermore, a comparable *cis*-element was recruited into PEPC in three grass species (Gupta, et al. 2020). Altogether, this shows that a *cis*-element repository already existing in the ancestral C_3_ species could be recruited for the C_4_ photosynthesis.

Interestingly, 23 of the 37 C_4_ReTFs exhibit higher transcript abundance in *Fso* than in *Fro*, suggesting that some modifications in the TFs had also occurred during the evolution. In this context, it has been also reported that the *cis*-elements from PPDK, CA (Kajala, et al. 2012) and NAD-ME (Brown, et al. 2011) from C_3_ species can achieve the C_4_-type cell specific accumulation of GUS in C_4_ species. Therefore, the TFs in C_4_ species might have gained an increased expression or altered their cellular expression pattern to use existing cis-elements and thereby gain new regulatory functions. For instance, it has been previously reported that nine TFs acquired shared MC- or BSC-preferential expression patterns in two divergent C_4_ lineages: *G. gynandra* and *Z. mays* (Aubry, et al. 2014).

It is worthy to note that, due to the absence of genome sequence and annotation for any Flaveria species so far, it remains difficult to estimate to which extent the *cis*-elements of the C_4_ orthologous genes in *Fso* were newly recruited and whether the C_4_-type expression patterns of the recruited C_4_TFs were established in the same species (*Fso*).

### Does Fra represent a dead-end for the C_4_ evolution?

The current notion of C_4_ evolution in the genus Flaveria is that the C_3_-C_4_ intermediates have two types, with *Fra* represents the type II C_3_-C_4_ intermediate, which shows more C_4_-ness than the type I C_3_-C_4_ species does (McKown and Dengler 2007; Vogan and Sage 2011). Given this notion, it is striking that *Fra* shows less C_4_-like C_4_GRN than *Fso* does, as reflected by the absence of a number of critical C_4_ genes in the overlapping C_4_GRN between *Fra* and *Ftr* (fig. 5*A*). Furthermore, in *Fra*, the genes in the MMC_4_ display no enriched GO functions in photosynthesis (fig. 4*D*). Therefore, it raises a question regarding whether *Fra* constitutes a transitional state from type I C_3_-C_4_ to C_4_ photosynthesis. On one hand, compared to type I C_3_-C_4_ intermediate, *Fra* indeed shows increased vein density (McKown and Dengler 2007), an increased cyclic electron transport capacity (Vogan and Sage 2011; Nakamura, et al. 2013), lower CO_2_ compensation point (Ku, et al. 1983; Nakamoto, et al. 1983; Rumpho, et al. 1984; Ku, et al. 1991) and higher expression levels for many C_4_ genes (Gowik, et al. 2011; Mallmann, et al. 2014). Besides, 42% of CO_2_ can initially be fixed into malate and aspartate in *Fra* (Rumpho, et al. 1984). However, on the other hand, *Fra* also shows C_3_ characteristics for a number of other C_4_ related features, such as CO_2_ assimilation rate (Ku, et al. 1991), δ^13^C (Gowik, et al. 2011) and the water using efficiency (Vogan and Sage 2011). The C_3_-level of the δ^13^C implies that there is no substantial increase in the C_4_ flux in the *Fra* compared to the C_3_ *Fro*.

However, we did find that four C_4_TF and PEPC1 co-regulations are conserved between *Fra* and *Ftr* (fig. 5*B*). Moreover, *Fra* shows C_4_ type MEM1 motif at its promoter region (Fig. 5*C*). All these are in line with the increased transcript abundance for PEPC1 in *Fra* compared with that of *Fro* and *Fso* (Gowik, et al. 2011; Mallmann, et al. 2014), which implies the increased role of PEPC in fixing CO_2_ in this species. However, the fixed CO_2_ might not be pumped to the BSC through the malate or aspartate as the typical C_4_ metabolic pathways do. Because if it is the case, an increase in δ^13^C would be expected, as for the typical C_4_ species (Farquhar, et al. 1989). Furthermore, neither NADP-MDH, which converts OAA to malate, nor AspAT, which converts OAA to aspartate, shows increased transcript abundance in *Fra* compared to the C_3_ *Fro* (supplementary fig. S2A-D; (Gowik, et al. 2011; Mallmann, et al. 2014). Therefore, the fixed CO_2_ in the form of OAA might be used to support metabolic reactions other than the C_4_ cycle.

If there is no increased C_4_ metabolic flux in *Fra*, so what might underlie the observed enhanced C_4_-ness in *Fra*? The high C_4_-ness of *Fra* most probably resulted from the increased C_2_ CCM. An increased PEPC1 function would result in a low CO_2_ concentration in the MC, which could increase the photorespiratory fluxes. Indeed, *Fra* has relatively higher expression levels for the photorespiratory genes if compared to both C_3_ and C_4_ species (Mallmann, et al. 2014). The increased photorespiratory flux in MC could enhance the glycine shuttle from the MC towards the BSC, where glycine would be converted to serine, leading to an accentuated release of the CO_2_ and ammonium in the BSC and consequently favorites the C_2_ CCM. The increased C_2_ CCM does not affect δ^13^C as much as in the case of C_4_. Furthermore, in *Fra*, the CO_2_ fixed by PEPC1 might be released into the cytosol through the TCA cycle. In another word, the primary metabolism of *Fra* may represent an alternative strategy to strengthen the CCM. In this regard, earlier studies showed that the intermediate species *Alloteropsis semialata* of the grass family (Lundgren, et al. 2016), C_3_-C_4_ intermediate species from Moricandia such as *M. arvensis* and *M. suffruticosa* (Schluter, et al. 2017), and C_3_-C_4_ species *Salsola divaricata* (Lauterbach, et al. 2017) displayed comparable values of δ^13^C to that of the C_3_ species. The metabolic status of the C_3_-C_4_ intermediate species deserves further detailed studies.

### Potential existence of a master regulator mediating the C_4_ photosynthesis

In the C_4_ species *Ftr*, many TFs were shown to be co-regulated with multiple C_4_ genes, *e.g*., two TFs (Gene24690 and Gene02109) are co-regulated with the same four C_4_ genes including PPDK-RP, BASS2, DiT2 and PPT1 (fig. 3*C* and Supplementary Material SF5). Furthermore, GRAS is the second most abundant TF family in the C_4_GRN of *Ftr*, with 21 GRAS TFs co-regulating 9 over 13 C_4_ genes (CA1, PEPC1, NADP-ME4, NADP-MDH, PPDK, PPDK-RP, DiT1, DiT2 and NHD1; fig. 3*F*). Given that the TFs of GRAS family are reported to also control the development of Kranz anatomy, such as Scarecrow (SCR), Shortroot (SHR) (Slewinski 2013; Wang, et al. 2013; Fouracre, et al. 2014) and Golden2-like (GLK) (Wang, et al. 2017), the TFs in the GRAS family might co-regulate the C_4_ metabolism and anatomy simultaneously. Another significant change in the C_4_TF network of *Ftr* compared to the three other Flaveria species is the recruitment of a large number of TFs for PEPC1 and PPDK-RP. This recruitment of TFs regulating PPDK-RP agrees with the crucial and prominent role for the PPDK-RP in maintaining a high efficient C_4_ metabolism in *Z. mays* (Hart, et al. 2011). Here we found that TFs from the GRAS family co-regulate these two enzymes (fig. 3*F*), again suggesting a possibility that members of the GRAS family TFs may represent candidate master regulators for C_4_ photosynthesis evolution in the genus Flaveria. Therefore, efforts are warranted to identify TFs co-regulating both C_4_ anatomical and metabolic features to support C_4_ crop engineering.

## Materials and Methods

### Plant materials and growth conditions

Seeds of *Fro* (C_3_) and *Fra* (C_3_-C_4_, type II) were provided by Prof. Peter Westhoff (Heinrich Heine University, Germany). Seeds of *Fso* (C_3_-C_4_, type I) and *Ftr* (C_4_) were obtained from Prof. Rowan Sage (University of Toronto, Canada). For low CO_2_ experiment, plants were grown in the growth chamber with a photosynthetic photon flux density (PPFD) controlled to be 200 μmol m^−2^ s^−1^, temperature 22 ±2 °C, 70% relative humidity (RH), and a photoperiod of 16 h light/8 h dark. The CO_2_ concentration used for the low CO_2_ treatment was 100 ppm and the control CO_2_ was around 380 ppm. Plants for ABA treatment were grown in the phytotron of Shanghai Institute of Plant Physiology and Ecology (SIPPE), Chines Academy of Sciences (CAS), the PPFD was fixed to 500 μmol m^−2^ s^−1^, and temperature 25 ±1 °C, 70% RH and the photoperiod was 16 h light/8h dark. The ABA was firstly dissolved in the distilled water to prepare a concentration of 40 μM ABA which was subsequently sprayed onto the mature leaves (2^nd^ or 3^rd^ pair of leaves counting from the top) of one-month old plants. During the ABA treatment, the mature leaves of control plants were sprayed only with distilled water (without ABA). For high light experiment, plants were grown under a phytotron in the Partner Institute of Computational Biology (PICB), CAS, with a PPFD of 1400 μmol m^−2^ s^−1^ (high light conditions) and that for the control condition was set to 500 μmol m^−2^ s^−1^. The high light condition was achieved by supplementing light with a lab-made light emitting diode (LED) light source. The light spectrum used in both control and high light conditions was as depicted in supplementary figure S6. For all experiments, plants were watered twice a week and fertilizer was weekly used after being dissolved to a concentration of 1‰ (w/w) (N: P: K = 20:20:20) to avoid plants growth limitation due to nutrition depletion and obtain healthy plants useful thereafter for our measurements.

### PacBio Iso-seq and next-generation sequencing of RNA

To obtain a better annotation for Flaveria genes and understand their potential biological functions, we sequenced full-length transcript sequences across the transcriptome of *Fra* using Iso-seq, which was used as a reference to annotate genes of the other three species (supplementary fig. S1). The RNA samples isolated from four different tissues, including leaf, stem, root and flower, of *Fra* were mixed and subjected to the sequencing process in order to obtain a blueprint of transcripts. For leaf samples, the younger fully expanded leaf from one-month old plants, which usually located on the 2^nd^ or the 3^rd^ pair of leaves counting from the top, was used. For the stem samples, the mid-segment of a stem between the root node and that of the leaf from one-month old plants was used. Concerning the root samples, the root hair of one-month old plants was used. The compacted soil on the roots was removed by floating and gently shaking root in water before samples for RNA were taken. Regarding the flower samples, three to five inflorescences from the same plant were sampled. Each sample was collected from three different plants. All samples were cut and immediately frozen in liquid nitrogen. Total RNA for each tissue was then isolated following the protocol of the PureLInk™ RNA kit (ThermoFisher Scientific, USA). The isolated RNA from the above mentioned four tissues was mixed in equal amount for the next step of analysis. The cDNA was synthesized using SMARTer^®^ PCR cDNA Synthesis Kit (Clontech, USA) which was then amplified using the KAPA HiFi PCR Kits (Roche, USA). The amplified products were classified into different fragments according to their sizes (1~2 k, 2~3 k or > 3 k) using BluePippin Size Selection (Sage Science, USA) which were subsequently utilized to construct a library separately through the SMRTbellTM Template Prep Kit 1.0 (PacBio, USA). Libraries were sequenced by the PacBio Sequel (PacBio, USA).

For the NGS of the RNA data, the younger fully expanded leaf obtained from 4-week (ABA, high light and 2-week low CO_2_), 6-week (4-week low CO_2_) and 6.5-month (6-month low CO_2_) old plants usually situated on the 2^nd^ or 3^rd^ pair of leaves counting started from the top was used for all the species studied herein. The chosen leaves were cut and immediately frozen into liquid nitrogen and stored thereafter at −80 °C until further processing. The total RNA was isolated according to the procedure described above. The NGS of RNA data was performed in the Illumina platform in the paired-end mode with a read length of 150 bp.

### Transcripts assembly, annotation and quantification

The Iso-seq long reads (LR) of *Fra* was corrected using proovread (Hackl, et al. 2014) with NGS of RNA-seq short reads (SR). In this process, the SR was mapped to LR and sequencing errors of LR were corrected by short-read-consensus approach (Hackl, et al. 2014). The rectified Iso-seq reads were then utilized to conduct transcript assembly applying IDP-*de novo* (Fu, et al. 2018). To define (or characterize) a reference on the gene scale, we used blast (V 2.2.31^+^) (Camacho, et al. 2009) to detect the transcript groups which have high similarity, *i.e*., those with an E-value threshold of 10^−10^ and the sequence identity threshold of 90%. The longest transcript in the transcript group was used as a reference gene. All the identified reference genes form together the gene reference dataset. The biological functions of *Fra* genes were annotated by seeking the orthologs in the database Uniprot (https://www.uniprot.org/). In this study, we used a combination of two datasets. The first represents the Uniprot reference clusters of Uniref50 (each entry represents a cluster of sequences with at least 50% sequence identity and 80% an overlap). For this first dataset, further details are available on the following website (https://www.uniprot.org/help/uniref). The second dataset constitutes the Swissprot (manually annotated and reviewed database). The Orthologs were predicted by using blast (V 2.2.31^+^) (Camacho, et al. 2009) with an E-value cutoff threshold of 10^−5^. The TFs were annotated by searching the orthologs of all plant TFs in the database PlantTFDB (http://planttfdb.cbi.pku.edu.cn/; (Riano-Pachon, et al. 2007) using blast (V 2.2.31^+^) (Camacho, et al. 2009) with an E-value threshold of 10^−5^ (supplementary fig. S1).

The transcript sequence of *Fro*, *Fso* and *Ftr* were *de novo* assembled based on SR. The *de novo* assembly was also performed for the SR in *Fra*. A Trinity (version 2.8.4) (Grabherr, et al. 2011) was used to perform the *de novo* assembly using default parameters except constraining the transcript length to be no less than 300 bp. The *de novo* assembled transcripts were annotated by searching the orthologs from *Fra* genes using blast (V 2.2.31^+^) (Camacho, et al. 2009) with an E-value threshold of 10^−5^.

The transcripts abundance was quantified as Transcripts per kilobase Per Million mapped reads (TPM) by applying the RSEM package (Li and Dewey 2011), where bowtie2 (version 2.3.4.3) (Langmead and Salzberg 2012) was used to map the SR to *de novo* assembled transcripts and the other parameters remained in default (supplementary figs. S2 and S3). The differentially expressed genes between, on one hand, the control and treated samples and, on the other hand, between the different species studied herein were calculated using edgeR package (Robinson, et al. 2010) with a *P-*value < 0.05 and FC > 1.5.

### qRT-PCR analysis

To verify the gene expressions determined based on the SR *de novo* assembly and confirm the accuracy of our computational analysis, a qRT-PCR analysis was conducted on the same RNA samples used for NGS of RNA data from leaf of six-week old plants including four-week treatment to low CO_2_ of 100 ppm. Between 0.2 and 0.5 μg RNA was used to reverse transcribe the first strand cDNA with Superscript II Reverse Transcriptase (TransGen Biotech, Beijing). The qRT-PCR mixture was prepared following the manufacturer’s instructions of UNICONTM qPCR SYBR Green Master Mix (YEASEN, Shanghai). Briefly, the mixture of cDNA, buffer and enzyme were pipetted into the Hard-Shell PCR 96-well plates (Bio-Rad, USA) and then covered by Microseal ‘B’ seal (Bio-Rad, USA). The qRT-PCR was run in the BIO-RAD CFX Connect-system (Bio-Rad, USA). Each PCR reaction volume of 20 μL contains 10 μL SYBR Green PCR Master Mix, 4 μL deionized H_2_O, 1 μL primers (0.5F + 0.5R) and 5 μL cDNA. The amplification reaction was initiated with a pre-denaturing step at 95 °C for 3 min followed by 39 cycles of denaturing at 95 °C for 10 s, annealing at 60 °C for 20 s, then extension at 72 °C for 30 s. The gene relative expression against the housekeeping gene, Actin7, was calculated as follows: 2^−ΔΔCT^ (Δ^CT^ = CT, gene of interest^−CT^, Actin7), as described previously by (Livak and Schmittgen 2001). Data was processed using Bio-Rad CFX Maestro software (Bio-Rad, USA). For each gene, two technical replicates and three biological replicates were performed. The primers used for qRT-PCR analysis are listed in supplementary table S3.

### Isolation of 5’ flanking sequence from PEPC1 gene of four species

The 5’ flanking sequence of the PEPC1 gene of the four Flaveria species (*Fro*, *Fso*, *Fra* and *Ftr*) was amplified from the total DNA through PCR reactions. The DNA was extracted following the protocol of TIANquick Midi Purificatioin kit (TIANGEN Biotech, Beijing). The primers used for the DNA amplification procedure with PCR are listed in supplementary table S4. Ultimately, the PCR products were sequenced in collaboration with Sangon Biotech Company (Shanghai, China).

### A GRN construction

The GRN was constructed with the CMIP software package (Zheng et al. 2016), which implements a path consistency algorithm based on the conditional mutual information (CMI) (Zhang, et al. 2012; Zheng, et al. 2016). The CMIP package uses a table of gene expressions as input. The method firstly computes zero-order network based on mutual information (MI), then eliminates the indirect relationship by considering the CMI, leading thereby to a first-order network. We also implemented the *P*-value determination in the CMIP package based on a permutation test. In other words, we shuffled 2000 times the expression level of each gene, generating thus 2000 null datasets. Thereafter, the *P*-value was defined as the proportion of CMI calculated from the null datasets greater than that calculated from the original dataset. We used a *P*-value of 0.001 as the cutoff during the genome-wide GRN reconstruction. The C_4_GRN was developed by only retaining the co-regulatory network of the C_4_ core metabolic genes and their positively co-regulated TFs (thereafter, C_4_TFs) from the genome-wide GRN.

In this study, special measures have been taken, thereafter, to ensure the development of a reliable and accurate GRN. In the original CMIP package, an adequate cutoff of gene-gene partial Pearson correlation coefficient (PCC) was automatically estimated by fitting the curve of interaction number in response to different cutoff based on an exponential function, and the cutoff was determined as the slopes intersection of the start and the end sections of the fitted curve (Zheng, et al. 2016). Given that we mainly focus on the C_4_GRN construction, we slightly modified this computational procedure. Specifically, we compared the relative abundance of all TF families (TF frequency) of the C_4_GRN under different PCC cutoffs, increasing from 0.1 to 0.9 with a step size of 0.1. Consequently, we found that the TF frequency maintains a high degree of similarity under different PCC cutoffs ranging from 0.1 to 0.7 (supplementary fig. S7). We calculated the similarity of TFs frequencies under two adjacent PCC cutoffs using Pearson correlation, and found the Pearson correlations are high (>0.95) from 0.1 to 0.7 but plummets from 0.7 to 0.8 (supplementary fig. S8). We thus selected 0.7 to be the PCC cutoff for the GRN construction. Interestingly, the number 0.7 represents the inflexion point on the curve, which (the curve) depicts the edge number versus different PCC cutoffs (supplementary fig. S9).

### Genome-wide GRN clustering and gene ontology enrichment analysis

In order to assess the potential implications of C_4_ genes and C_4_TFs in the different species such as their biological functions and/or the molecular relevance of their concurrent co-regulations, the genome-wide GRN was clustered into different modules. The latters (modules) enriched in C_4_TFs or those including the maximum C_4_ genes were selected for the functional enrichment analysis. Herein, we used a Markov Cluster Algorithm (MCL) package (version 14-137; (Dongen 2008) to detect the modules in the GRN. The MCL is an unsupervised model, where the number of clusters cannot be determined on the fly. The parameter “I” (inflation) represents the main handle affecting the cluster granularity and influences the cluster number and clustering efficiency that the algorithm produces (Enright, et al. 2002). The clustering efficiency was automatically calculated in the program by considering two factors. These are; the density of all clusters and the percentage of lost interactions. In order to determine a proper value for the “I” parameter used in MCL package, we examined the clustering efficiencies and cluster numbers for five different cutoffs of “I” including 1.4, 2, 4, 5, and 6. Then, we have chosen the ideal value for “I” which had the maximum clustering efficiency for a minimum clusters number. We found that all the four species investigated herein show the best “I” to be 2 (supplementary fig. S10A), with a clustering efficiency being 0.21, 0.2, 0.16 and 0.22 for *Fro*, *Fso*, *Fra* and *Ftr*, respectively.

Subsequently, we determined the modules enriched in the C_4_TFs (MET) using the *Fisher*’s test computed in-home R script. The calculated *P-*values from the *Fisher’s* test were adjusted using the Benjamini-Hochberg (BH) method (Benjamini and Hochberg 1995) integrated in R software (version 3.0.2). A threshold of 0.001 for the adjusted *P-*values was used. Besides, the modules including the maximum of C_4_ orthologous gene were termed as MMC_4_. To investigate the potential involvements of MET and MMC_4_ in certain biological functions, we calculated the over-represented gene ontology (GO) for each MET and MMC_4_ using *Fisher*’s test with BH adjusted *P* < 0.001, where the genes in the genome-wide GRN were used as a background.

### Accession number

The RNA-seq and Iso-seq data were submitted to Gene Expression Omnibus (GEO) in the Nation Center for Biotechnology Information (NCBI) database under the following accession number: GSE143470.

## Author contributions

XGZ, GC, MJL and JE designed the project and wrote the paper, MJL and FC did bioinformatics analysis, MJL did the qRT-PCR and JE did the RNA-isolation.

## Acknowledgements

The authors thank and appreciate Profs. Rowan F. Sage and Peter Westhoff for sharing with us Flaveria materials. We thank Prof. Haiyang Hu and Mingnan Qu for their fruitful discussions and suggestions. Authors thank also Fengfeng Miao, Qingfeng Song and Xinyu Liu for their assistance in maintaining the growth chambers conditions properly and according to the recorded instructions. This work was supported by National Science Foundation of China, NSFC (31701139, 31870214, 31500988), Strategic Priority Research Program of the Chinese Academy of Sciences, CAS (Grant No. XDB27020105), National Research and Development Program of Ministry of Science and Technology of China, MSTC (2019YFA0904600).

## Supplementary Material

SF1: supplementary figures and tables

SF2: Functional annotations of *Fra* genes

SF3: RNA-seq mapping statistics

SF4: Gene expression in TPM

SF5: C_4_GRN tables.

SF6: C_4_TFs enriched GOs

SF7: PEPC1 promoter sequences of the four Flaveria species

SF8: Functional annotations of the 37 C_4_TFs overlapped between *Fso* and *Ftr*

## References

Akyildiz M, Gowik U, Engelmann S, Koczor M, Streubel M, Westhoff P. 2007. Evolution and function of a cis-regulatory module for mesophyll-specific gene expression in the c_4_ dicot flaveria trinervia. Plant Cell 19:3391–3402.

Aldous SH, Weise SE, Sharkey TD, Waldera-Lupa DM, Stuhler K, Mallmann J, Groth G, Gowik U, Westhoff P, Arsova B. 2014. Evolution of the phosphoenolpyruvate carboxylase protein kinase family in c_3_ and c_4_ flaveria spp. Plant Physiol 165:1076–1091.

Aubry S, Kelly S, Kumpers BM, Smith-Unna RD, Hibberd JM. 2014. Deep evolutionary comparison of gene expression identifies parallel recruitment of trans-factors in two independent origins of c4 photosynthesis. PLoS Genet 10:e1004365.

Benjamini Y, Hochberg Y. 1995. Controlling the false discovery rate - a practical and powerful approach to multiple testing. Journal of the Royal Statistical Society Series B-Statistical Methodology 57:289–300.

Brown NJ, Newell CA, Stanley S, Chen JE, Perrin AJ, Kajala K, Hibberd JM. 2011. Independent and parallel recruitment of preexisting mechanisms underlying c4 photosynthesis. Science 331:1436–1439.

Burgess SJ, Granero-Moya I, Grange-Guermente MJ, Boursnell C, Terry MJ, Hibberd JM. 2016. Ancestral light and chloroplast regulation form the foundations for c_4_ gene expression. Nat Plants 2:16161.

Burgess SJ, Reyna-Llorens I, Stevenson SR, Singh P, Jaeger K, Hibberd JM. 2019. Genome-wide transcription factor binding in leaves from c3 and c4 grasses. Plant Cell 31:2297–2314.

Camacho C, Coulouris G, Avagyan V, Ma N, Papadopoulos J, Bealer K, Madden TL. 2009. Blast plus: Architecture and applications. BMC Bioinformatics 10.

Cheng SH, Moore BD, Edwards GE, Ku MSB. 1988. Photosynthesis in flaveria-brownii, a c-4-like species - leaf anatomy, characteristics of co2 exchange, compartmentation of photosynthetic enzymes, and metabolism of (co2)-c-14. Plant Physiol 87:867–873.

Christin PA, Boxall SF, Gregory R, Edwards EJ, Hartwell J, Osborne CP. 2013. Parallel recruitment of multiple genes into c4 photosynthesis. Genome Biology and Evolution 5:2174–2187.

Christin PA, Osborne CP, Sage RF, Arakaki M, Edwards EJ. 2011. C4 eudicots are not younger than c4 monocots. Journal of Experimental Botany 62:3171–3181.

Christin PA, Petitpierre B, Salamin N, Buchi L, Besnard G. 2009. Evolution of c_4_ phosphoenolpyruvate carboxykinase in grasses, from genotype to phenotype. Molecular Biology and Evolution 26:357–365.

Dmoore B, Ku MSB, Edwards GE. 1989. Expression of c_4_-like photosynthesis in several species of flaveria. Plant Cell and Environment 12:541–549.

Dongen SV. 2008. Graph clustering via a discrete uncoupling process. SIAM Journal on Matrix Analysis and Applications 30:121–141.

Duarte KE, de Souza WR, Santiago TR, Sampaio BL, Ribeiro AP, Cotta MG, da Cunha B, Marraccini PRR, Kobayashi AK, Molinari HBC. 2019. Identification and characterization of core abscisic acid (aba) signaling components and their gene expression profile in response to abiotic stresses in setaria viridis. Sci Rep 9:4028.

Edwards GE, Ku MSB. 1987. Biochemistry of c_3_-c_4_ intermediates. In: Hatch md, boardman nk, editors. The biochemistry of plants. New York: Academic Press:275–325.

Enright AJ, Van Dongen S, Ouzounis CA. 2002. An efficient algorithm for large-scale detection of protein families. Nucleic Acids Res 30:1575–1584.

Farquhar GD, Ehleringer JR, Hubick KT. 1989. Carbon isotope discrimination and photosynthesis. Annual Review of Plant Physiology and Plant Molecular Biology 40:503–537.

Fischer E, Raschke K, Stitt M. 1986. Effects of abscisic-acid on photosynthesis in whole leaves - changes in co_2_ assimilation, levels of carbon-reduction-cycle intermediates, and activity of ribulose-1,5-bisphosphate carboxylase. Planta 169:536–545.

Fouracre JP, Ando S, Langdale JA. 2014. Cracking the kranz enigma with systems biology. Journal of Experimental Botany 65:3327–3339.

Friedman N. 2004. Inferring cellular networks using probabilistic graphical models. Science 303:799–805.

Fu S, Ma Y, Yao H, Xu Z, Chen S, Song J, Au KF. 2018. Idp-denovo: De novo transcriptome assembly and isoform annotation by hybrid sequencing. Bioinformatics 34:2168–2176.

Fukayama H, Tsuchida H, Agarie S, Nomura M, Onodera H, Ono K, Lee BH, Hirose S, Toki S, Ku MS, et al. 2001. Significant accumulation of c(4)-specific pyruvate, orthophosphate dikinase in a c(3) plant, rice. Plant Physiology 127:1136–1146.

Gardner TS, di Bernardo D, Lorenz D, Collins JJ. 2003. Inferring genetic networks and identifying compound mode of action via expression profiling. Science 301:102–105.

Gorska AM, Gouveia P, Borba AR, Zimmermann A, Serra TS, Lourenco TF, Margarida Oliveira M, Peterhansel C, Saibo NJM. 2019. Zmbhlh80 and zmbhlh90 transcription factors act antagonistically and contribute to regulate pepc1 cell-specific gene expression in maize. Plant J 99:270–285.

Gowik U, Brautigam A, Weber KL, Weber AP, Westhoff P. 2011. Evolution of c4 photosynthesis in the genus flaveria: How many and which genes does it take to make c4? the Plant Cell 23:2087–2105.

Gowik U, Burscheidt J, Akyildiz M, Schlue U, Koczor M, Streubel M, Westhoff P. 2004. Cis-regulatory elements for mesophyll-specific gene expression in the c_4_ plant flaveria trinervia, the promoter of the c_4_ phosphoenolpyruvate carboxylase gene. Plant Cell 16:1077–1090.

Gowik U, Schulze S, Saladie M, Rolland V, Tanz SK, Westhoff P, Ludwig M. 2017. A mem1-like motif directs mesophyll cell-specific expression of the gene encoding the c4 carbonic anhydrase in flaveria. Journal of Experimental Botany 68:311–320.

Grabherr MG, Haas BJ, Yassour M, Levin JZ, Thompson DA, Amit I, Adiconis X, Fan L, Raychowdhury R, Zeng QD, et al. 2011. Full-length transcriptome assembly from rna-seq data without a reference genome. Nat Biotechnol 29:644–U130.

Gupta SD, Levey M, Schulze S, Karki S, Emmerling J, Streubel M, Gowik U, Paul Quick W, Westhoff P. 2020. The c_4_ ppc promoters of many c_4_ grass species share a common regulatory mechanism for gene expression in the mesophyll cell. Plant J 101:204–216.

Hackl T, Hedrich R, Schultz J, Forster F. 2014. Proovread: Large-scale high-accuracy pacbio correction through iterative short read consensus. Bioinformatics 30:3004–3011.

Hart Y, Mayo AE, Milo R, Alon U. 2011. Robust control of pep formation rate in the carbon fixation pathway of c_4_ plants by a bi-functional enzyme. BMC Syst Biol 5:171.

Hatch MD. 1987. C_4_ photosynthesis - a unique blend of modified biochemistry, anatomy and ultrastructure. Biochimica et Biophysica Acta 895:81–106.

Hibberd JM, Covshoff S. 2010. The regulation of gene expression required for c_4_ photosynthesis. Annu Rev Plant Biol 61:181–207.

Hibberd JM, Sheehy JE, Langdale JA. 2008. Using c_4_ photosynthesis to increase the yield of rice-rationale and feasibility. Curr Opin Plant Biol 11:228–231.

Jin J, Tian F, Yang DC, Meng YQ, Kong L, Luo J, Gao G. 2017. Planttfdb 4.0: Toward a central hub for transcription factors and regulatory interactions in plants. Nucleic Acids Res 45:D1040–D1045.

Kajala K, Brown NJ, Williams BP, Borrill P, Taylor LE, Hibberd JM. 2012. Multiple arabidopsis genes primed for recruitment into c_4_ photosynthesis. Plant Journal 69:47–56.

Ku MS, Agarie S, Nomura M, Fukayama H, Tsuchida H, Ono K, Hirose S, Toki S, Miyao M, Matsuoka M. 1999. High-level, expression of maize phosphoenolpyruvate carboxylase in transgenic rice plants. Nat Biotechnol 17:76–80.

Ku MSB, Nakamoto H, Monson RK, Littlejohn RO, Fisher DB, Edwards GE. 1983. Photosynthetic characteristics of flaveria species intermediate between c-3 and c-4 plants. Plant Physiol 72:43–43.

Ku MSB, Wu JR, Dai ZY, Scott RA, Chu C, Edwards GE. 1991. Photosynthetic and photorespiratory characteristics of flaveria species. Plant Physiol 96:518–528.

Langmead B, Salzberg SL. 2012. Fast gapped-read alignment with bowtie 2. Nature Methods 9:357–U354.

Lauterbach M, Schmidt H, Billakurthi K, Hankeln T, Westhoff P, Gowik U, Kadereit G. 2017. De novo transcriptome assembly and comparison of c_3_, c_3_-c_4_, and c_4_ species of tribe salsoleae (chenopodiaceae). Front Plant Sci 8:1939.

Li B, Dewey CN. 2011. Rsem: Accurate transcript quantification from rna-seq data with or without a reference genome. BMC Bioinformatics 12:323.

Li Y, Xu J, Haq NU, Zhang H, Zhu XG. 2014. Was low co2 a driving force of c4 evolution: Arabidopsis responses to long-term low co2 stress. Journal of Experimental Botany 65:3657–3667.

Livak KJ, Schmittgen TD. 2001. Analysis of relative gene expression data using real-time quantitative pcr and the 2(t)(-delta delta c) method. Methods 25:402–408.

Lundgren MR, Christin PA. 2017. Despite phylogenetic effects, c-3-c-4 lineages bridge the ecological gap to c-4 photosynthesis. Journal of Experimental Botany 68:241–254.

Lundgren MR, Christin PA, Escobar EG, Ripley BS, Besnard G, Long CM, Hattersley PW, Ellis RP, Leegood RC, Osborne CP. 2016. Evolutionary implications of c3-c4 intermediates in the grass alloteropsis semialata. Plant Cell Environ 39:1874–1885.

Lyu MJ, Gowik U, Kelly S, Covshoff S, Mallmann J, Westhoff P, Hibberd JM, Stata M, Sage RF, Lu H, et al. 2015. Rna-seq based phylogeny recapitulates previous phylogeny of the genus flaveria (asteraceae) with some modifications. BMC Evol Biol 15:116.

Mallmann J, Heckmann D, Brautigam A, Lercher MJ, Weber AP, Westhoff P, Gowik U. 2014. The role of photorespiration during the evolution of c4 photosynthesis in the genus flaveria. eLife 3:e02478.

McKown AD, Dengler NG. 2007. Key innovations in the evolution of kranz anatomy and c4 vein pattern in flaveria (asteraceae). Am J Bot 94:382–399.

McKown AD, Moncalvo J-M, Dengler NG. 2005. Phylogeny of flaveria (asteraceae) and inference of c4 photosynthesis evolution. American Journal of Botany 92:1911–1928.

Moreno-Villena JJ, Dunning LT, Osborne CP, Christin PA. 2018. Highly expressed genes are preferentially co-opted for c4 photosynthesis. Molecular Biology and Evolution 35:94–106.

Nakamoto H, Ku MSB, Edwards GE. 1983. Photosynthetic characteristics of c3-c4 intermediate flaveria species.2. Kinetic-properties of phosphoenolpyruvate carboxylase from c-3, c-4 and c3-c4 intermediate species. Plant and Cell Physiology 24:1387–1393.

Nakamura N, Iwano M, Havaux M, Yokota A, Munekage YN. 2013. Promotion of cyclic electron transport around photosystem i during the evolution of nadp-malic enzyme-type c4 photosynthesis in the genus flaveria. New Phytol 199:832–842.

Nomura M, Sentoku N, Nishimura A, Lin JH, Honda C, Taniguchi M, Ishida Y, Ohta S, Komari T, Miyao-Tokutomi M, et al. 2000. The evolution of c4 plants: Acquisition of cis-regulatory sequences in the promoter of c4-type pyruvate, orthophosphate dikinase gene. Plant Journal 22:211–221.

Powell AM. 1978. Systematics of flaveria (flaveriinae asteraceae). Annals of the Missouri Botanical Garden 65:590–636.

Reyna-Llorens I, Burgess SJ, Reeves G, Singh P, Stevenson SR, Williams BP, Stanley S, Hibberd JM. 2018. Ancient duons may underpin spatial patterning of gene expression in c_4_ leaves. Proc Natl Acad Sci U S A 115:1931–1936.

Riano-Pachon DM, Ruzicic S, Dreyer I, Mueller-Roeber B. 2007. Plntfdb: An integrative plant transcription factor database. BMC Bioinformatics 8.

Robinson MD, McCarthy DJ, Smyth GK. 2010. Edger: A bioconductor package for differential expression analysis of digital gene expression data. Bioinformatics 26:139–140.

Rumpho ME, Ku MSB, Cheng SH, Edwards GE. 1984. Photosynthetic characteristics of c3-c4 intermediate flaveria species.3. Reduction of photorespiration by a limited-c4 pathway of photosynthesis in flaveria-ramosissima. Plant Physiol 75:993–996.

Sage RF. 2001. Environmental and evolutionary preconditions for the origin and diversification of the c4 photosynthetic syndrome. Plant Biol (Stuttg) 3:202–213.

Sage RF. 2004. The evolution of c4 photosynthesis. New Phytologist:341–370.

Sage RF, Sage TL, Kocacinar F. 2012. Photorespiration and evolution of c_4_ photosynthesis. Annual Review of Plant Biologist 63:19–47.

Sage RF, Zhu XG. 2011. Exploiting the engine of c4 photosynthesis. Journal of Experimental Botany 62:2989–3000.

Schluter U, Brautigam A, Gowik U, Melzer M, Christin PA, Kurz S, Mettler-Altmann T, Weber AP. 2017. Photosynthesis in c3-c4 intermediate moricandia species. Journal of Experimental Botany 68:191–206.

Schluter U, Weber APM. 2020. Regulation and evolution of c_4_ photosynthesis. Annu Rev Plant Biol 71:183–215.

Slewinski TL. 2013. Using evolution as a guide to engineer kranz-type c4 photosynthesis. Front Plant Sci 4.

Thompson D, Regev A, Roy S. 2015. Comparative analysis of gene regulatory networks: From network reconstruction to evolution. Annu Rev Cell Dev Biol 31:399–428.

Ubierna N, Sun W, Kramer DM, Cousins AB. 2013. The efficiency of c4 photosynthesis under low light conditions in zea mays, miscanthus x giganteus and flaveria bidentis. Plant Cell Environ 36:365–381.

Ueno O. 2001. Environmental regulation of c(3) and c(4) differentiation in the amphibious sedge eleocharis vivipara. Plant Physiol 127:1524–1532.

Vogan PJ, Sage RF. 2011. Water-use efficiency and nitrogen-use efficiency of c_3_-c_4_ intermediate species of flaveria juss. (asteraceae). Plant Cell Environ 34:1415–1430.

Wang P, Kelly S, Fouracre JP, Langdale JA. 2013. Genome-wide transcript analysis of early maize leaf development reveals gene cohorts associated with the differentiation of c4 kranz anatomy. Plant J 75:656–670.

Wang P, Khoshravesh R, Karki S, Tapia R, Balahadia CP, Bandyopadhyay A, Quick WP, Furbank R, Sage TL, Langdale JA. 2017. Re-creation of a key step in the evolutionary switch from c3 to c4 leaf anatomy. Curr Biol 27:3278–3287 e3276.

Williams BP, Aubry S, Hibberd JM. 2012. Molecular evolution of genes recruited into c4 photosynthesis. Trends Plant Sci 17:213–220.

Williams BP, Burgess SJ, Reyna-Llorens I, Knerova J, Aubry S, Stanley S, Hibberd JM. 2016. An untranslated cis-element regulates the accumulation of multiple c4 enzymes in gynandropsis gynandra mesophyll cells. Plant Cell 28:454–465.

Xu J, Brautigam A, Weber AP, Zhu XG. 2016. Systems analysis of cis-regulatory motifs in c_4_ photosynthesis genes using maize and rice leaf transcriptomic data during a process of de-etiolation. Journal of Experimental Botany 67:5105–5117.

Zhang XJ, Zhao XM, He K, Lu L, Cao YW, Liu JD, Hao JK, Liu ZP, Chen LN. 2012. Inferring gene regulatory networks from gene expression data by path consistency algorithm based on conditional mutual information. Bioinformatics 28:98–104.

Zheng G, Xu Y, Zhang X, Liu ZP, Wang Z, Chen L, Zhu XG. 2016. Cmip: A software package capable of reconstructing genome-wide regulatory networks using gene expression data. BMC Bioinformatics 17:535.

Zhu XG, Long SP, Ort DR. 2008. What is the maximum efficiency with which photosynthesis can convert solar energy into biomass? Curr Opin Biotechnol 19:153–159.

