## Supplemental File 1 for "Evolution of co-regulatory network of C_4_ metabolic genes and TFs in the genus Flaveria: go anear or away in the intermediate species?"

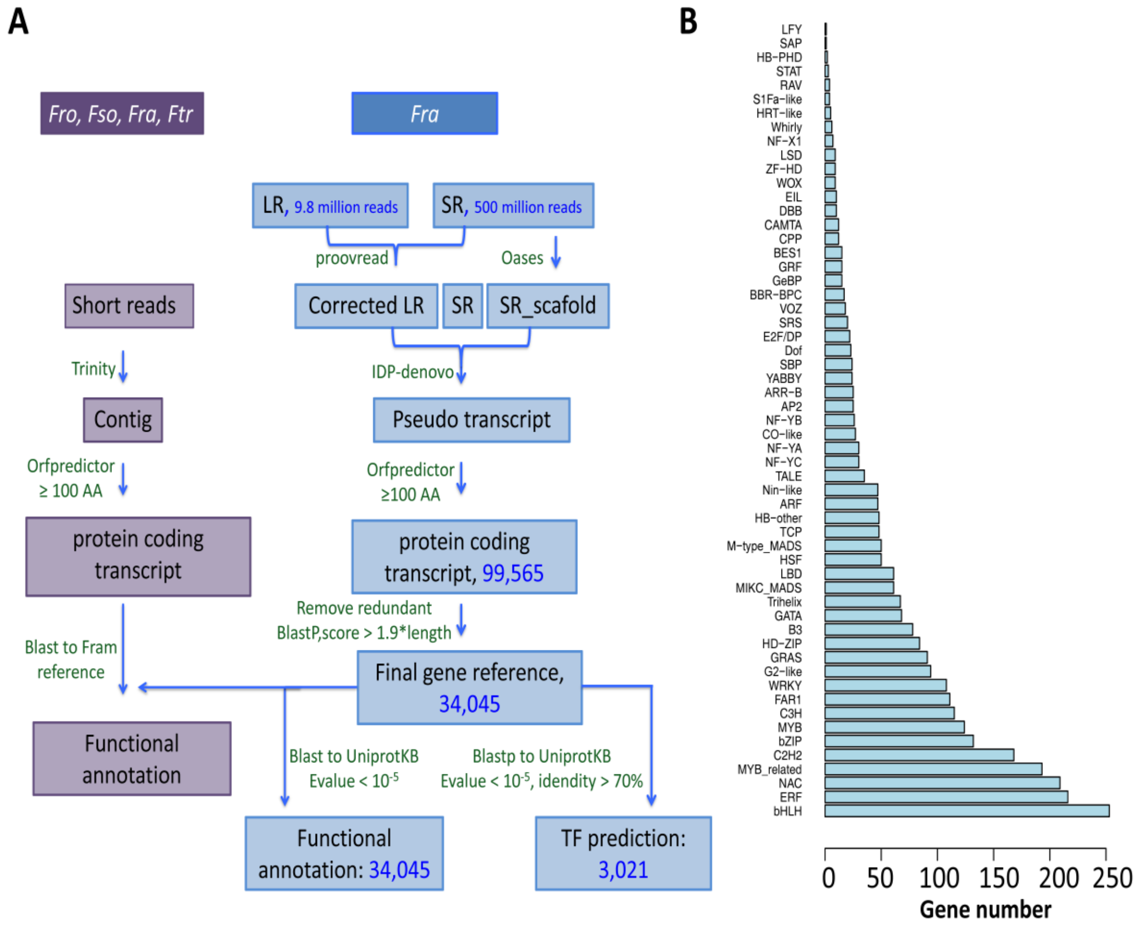


**Figure S1. Workflow of transcript assembly and annotation for four Flaveria species**

(A) Diagram of RNA-seq analysis process. The sequential events (steps) on right branch of the chart (in blue) explain the analysis pipeline of the PacBio Iso-seq data of *Fra*, and on the left part (in purple) shows the next generation RNA-seq data of all the four species (*Fro*, *Fso*, *Fra* and *Ftr*). (B) A bar plot showing the distribution of TF families in *Fra*. The values on the horizontal axis represent the genes number in each TF family. The TF families in the top of the pyramid show lesser than 50 genes by family. However, moving down to the bottom of the pyramid the genes number per family increases till it reaches around 250 genes in BHLH. Abbreviations: *Fro*, *F. robusta*; *Fso*, *F. sonorensis*; *Fra*, *F. ramosissima*; *Ftr*, *F. trinervia*.

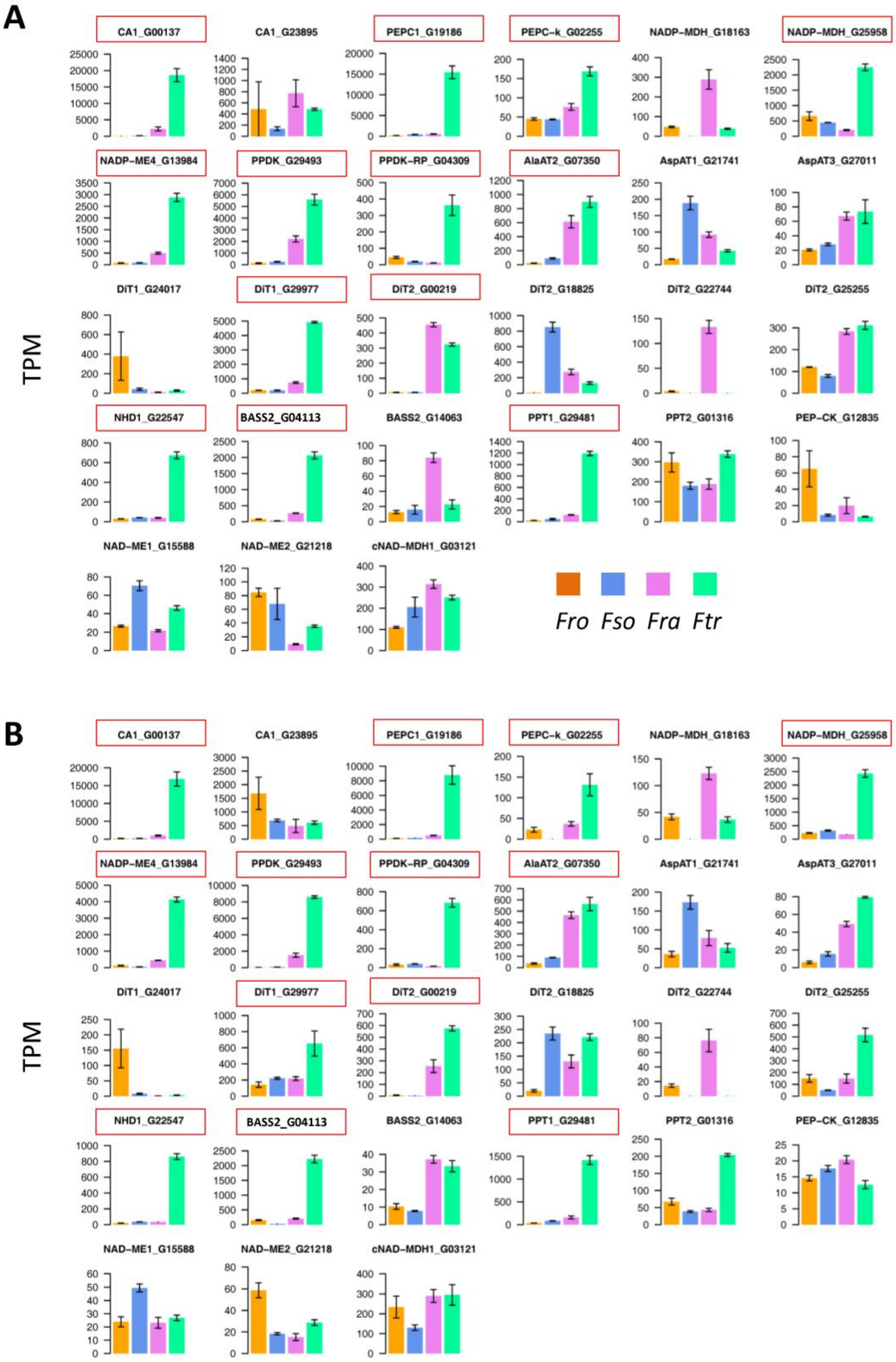
A
CA1_G00137
CA1_G23895
PEPC1_G19186
PEPC-k_G02255
NADP-MDH_G18163
NADP-MDH_G25958
NADP-ME4_G13984
PPDK_G29493
PPDK-RP_G04309
AlaAT2_G07350
AspAT1_G21741
AspAT3_G27011
DiT1_G24017
DiT1_G29977
DiT2_G00219
DiT2_G18825
DiT2_G22744
DiT2_G25255
NHD1_G22547
BASS2_G04113
BASS2_G14063
PPT1_G29481
PPT2_G01316
PEP-CK_G12835
NAD-ME1_G15588
NAD-ME2_G21218
cNAD-MDH1_G03121
TPM
Fro Fso Fra Ftr
B
CA1_G00137
CA1_G23895
PEPC1_G19186
PEPC-k_G02255
NADP-MDH_G18163
NADP-MDH_G25958
NADP-ME4_G13984
PPDK_G29493
PPDK-RP_G04309
AlaAT2_G07350
AspAT1_G21741
AspAT3_G27011
DiT1_G24017
DiT1_G29977
DiT2_G00219
DiT2_G18825
DiT2_G22744
DiT2_G25255
NHD1_G22547
BASS2_G04113
BASS2_G14063
PPT1_G29481
PPT2_G01316
PEP-CK_G12835
NAD-ME1_G15588
NAD-ME2_G21218
cNAD-MDH1_G03121
TPM

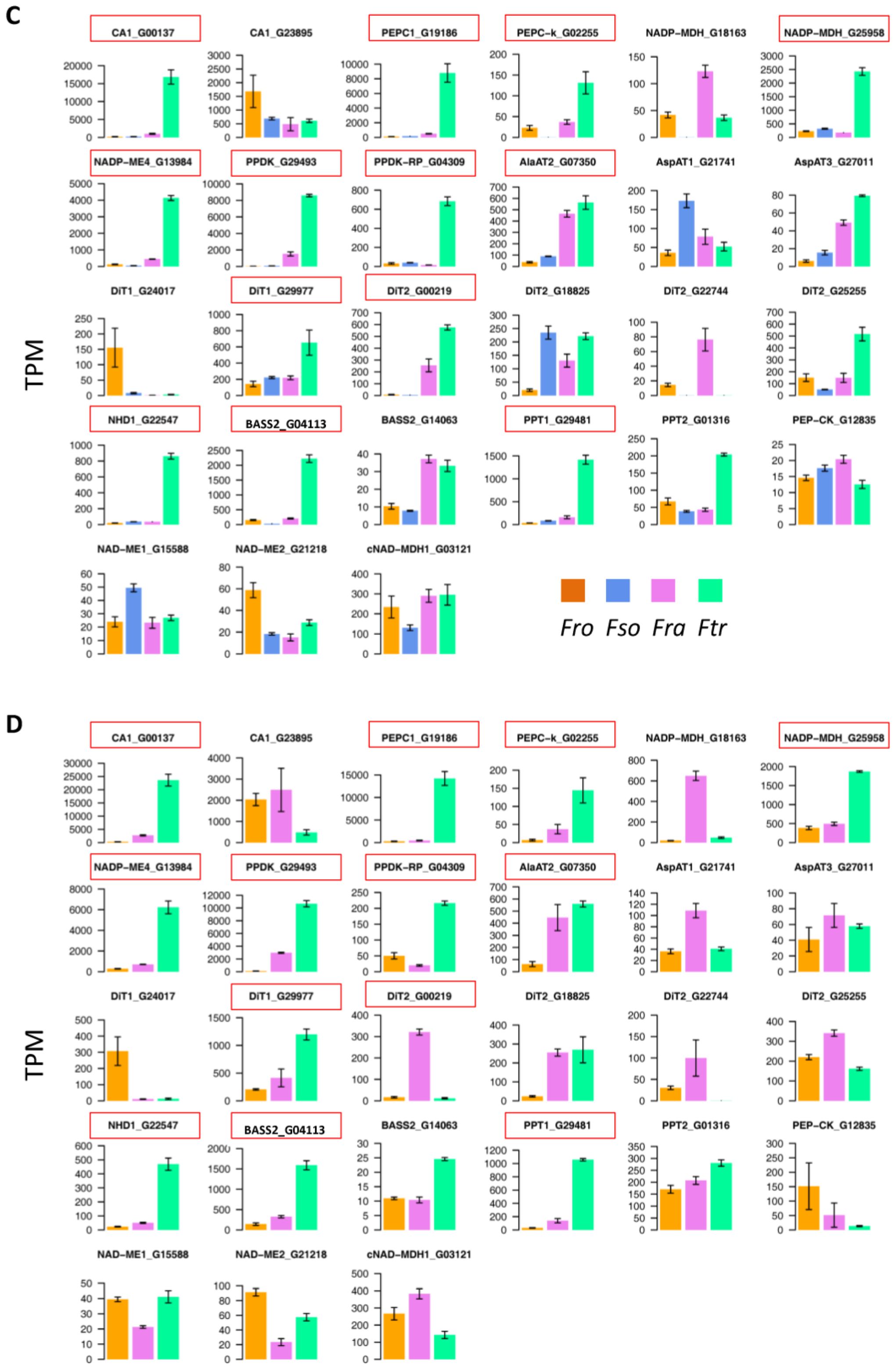


**Figure S2. Determination of the $C_4$ version of $C_4$ genes in Flaveria species**

The transcripts abundance of $C_4$ genes in four Flaveria species evaluated on plants exposed for two weeks (A) or four weeks to low $CO_2$ of 100 ppm (B), ABA for 3 hours (C) or high light for 2 weeks (D). *Fso* data were absent for the high light treatment (panel D). The $C_4$ version of $C_4$ genes (enclosed in a red rectangle) was determined based on two criteria. Firstly, it should show relatively higher transcript abundance than the other paralogs. Secondly, it must display higher transcript abundance in $C_4$ species than in $C_3$ species. Flaveria species abbreviations are as displayed in Fig. S1. TPM stands for **T**ranscripts per kilobase **P**er **M**illion mapped reads.

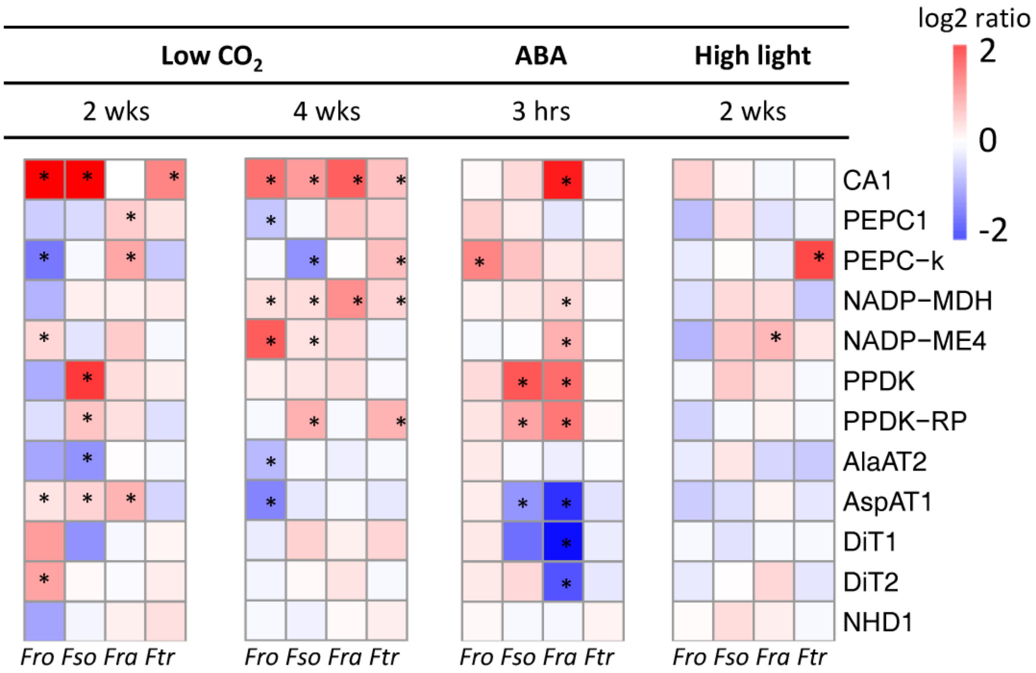


**Figures S3. Response of $C_4$ version of $C_4$ genes to different stress treatments**
Heatmaps show the log2 transformed fold change (FC) of transcript abundance for each treatment to that of control condition (FC = treated/control) in each experiment. Red represents increased TPM under treatment compared to control and blue represents decreased TPM compared to control. The star reveals the significant changes between treated and control samples at a *P*-value < 0.05 and FC > 1.5. TPM stands for **T**ranscripts per kilobase **P**er **M**illion mapped reads. hrs and wks stand hours and weeks, respectively. Flaveria species abbreviations are as displayed in Fig. S1

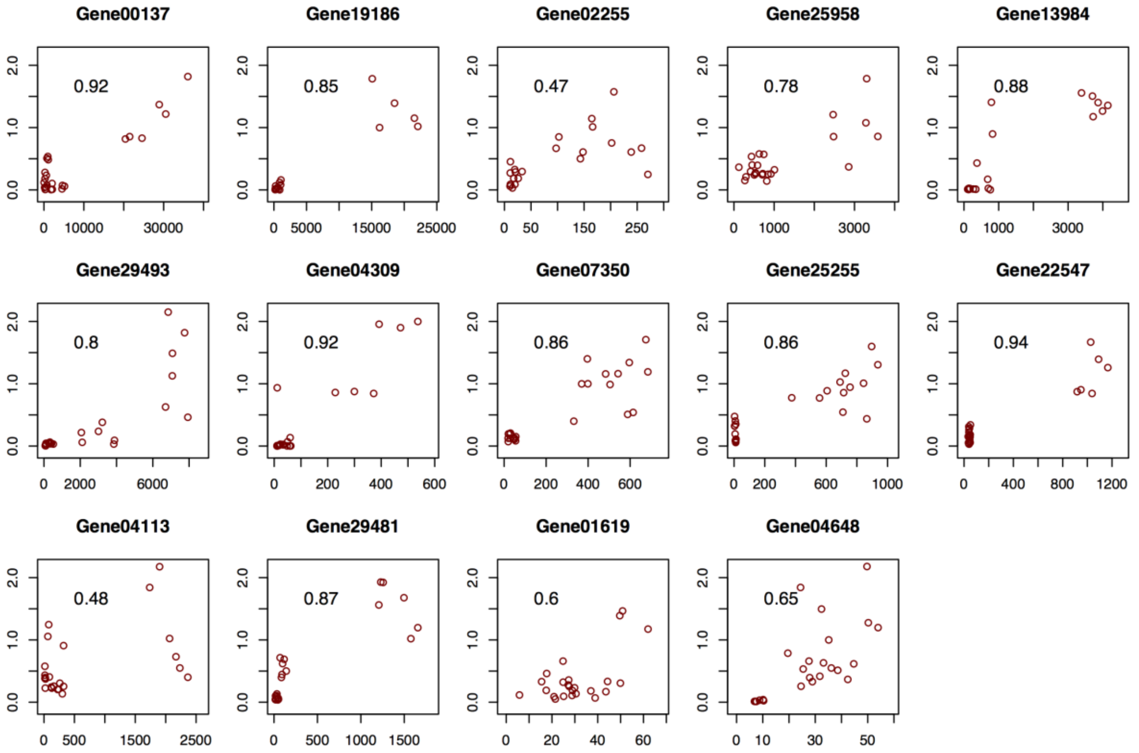


**Figures S4. Comparison of the quantification between RNA-seq data and qRT-PCR analysis**

The expression levels of 14 genes from RNA-seq were compared with their respective expression levels assessed through qRT-PCR analysis. The TMP (**T**ranscripts per kilobase **P**er **M**illion mapped reads) from RNA-seq (X-axis) was plotted against the relative expression from qRT-PCR (Y-axis). The RNA-seq data were obtained from plants treated with 100 ppm $CO_2$ experiment for four weeks. 24 samples were used from four Flaveria species. Each data point represents an individual sample. The values on the top left of each plot represent the Pearson correlation coefficient (PCC) between the abundance of 24 samples by RNA-seq and that by qRT-PCR. The PCC between RNA-seq data and the qRT-PCR analysis for the 14 genes was at least 0.47.

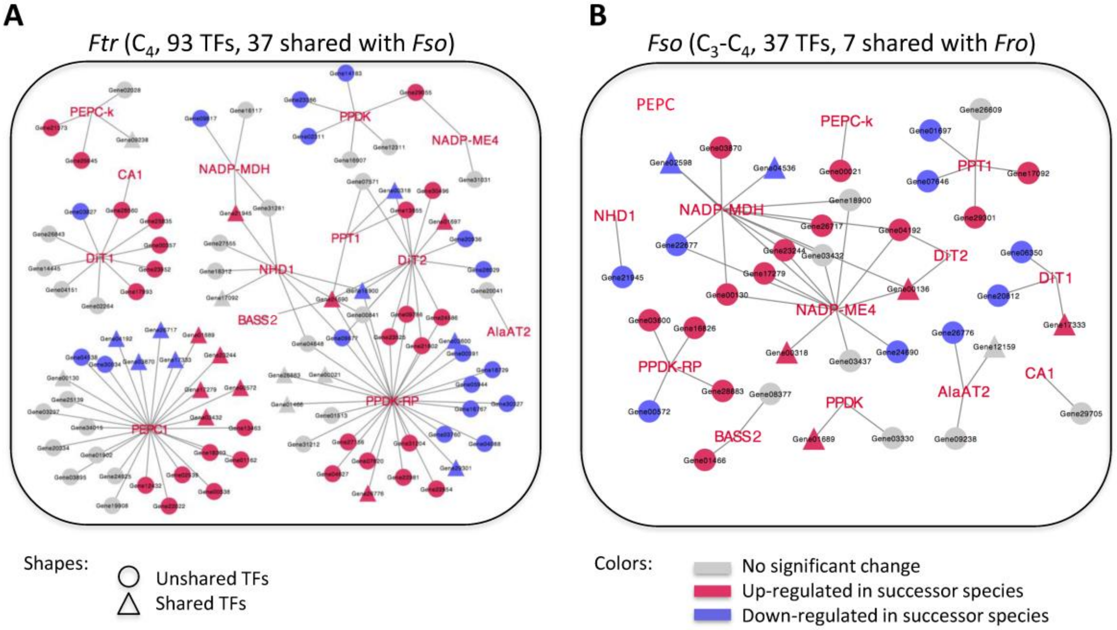


**Figure S5. The evolution of $C_4$GRN towards a $C_4$ photosynthesis stage**

(A) In *Ftr*, 93 $C_4$GRN TFs show higher transcript abundance than *Fro* (*P* < 0.05 and FC > 0.5*)*, termed as $C_4$-required TFs ($C_4$ReTFs), which co-regulate all the 13 $C_4$ core metabolic genes. (B) 37 of $C_4$ReTFs are present in the $C_4$GRN of *Fso*, co-regulated with all $C_4$ core metabolic genes except PEPC1. TFs are shown in circles and triangles, with triangles presenting TFs shared with either *Fso* (panel A) or *Fro* (panel B). Red, blue, and grey colors represent TF showing higher, lower and similar transcript abundance compared to *Fso* (A) or *Fro* (B) (edgeR, P < 0.01 and FC > 1.5). Abbreviations of the Flaveria species and the enzymes are the same as those shown in Figs. 1 and 2, respectively. FC stands for fold change.

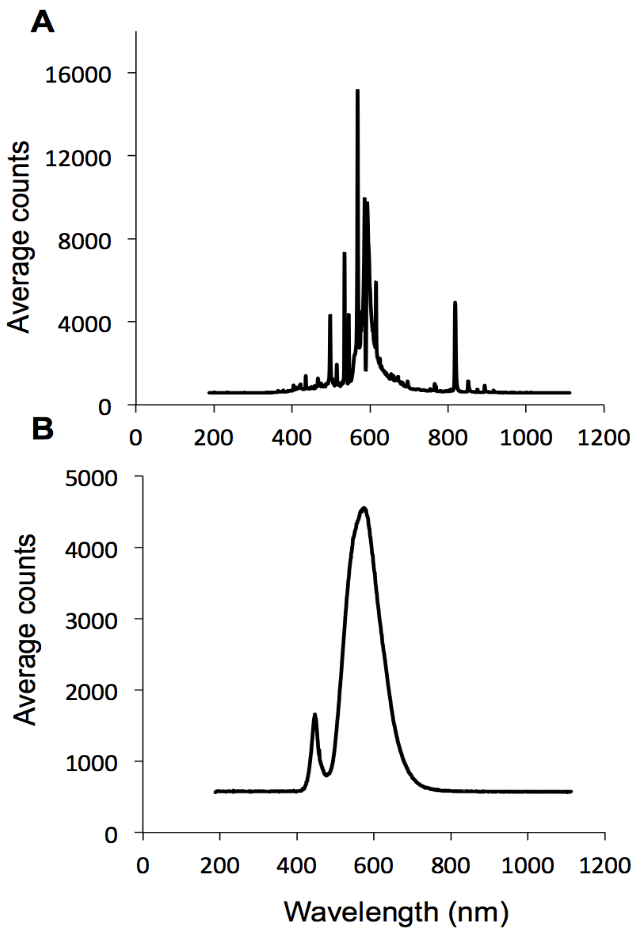


**Figure S6. Light spectrum used for high light experiment**
Spectral scans of the incident photosynthetic photo flux density (PPFD) for control condition light, 500 μmol $m^{-2}$ $s^{-1}$ (A) and high light treatment, 1400 μmol $m^{-2}$ $s^{-1}$ (B). High light condition was set up by a lab-made light emitting diode (LED) light source.

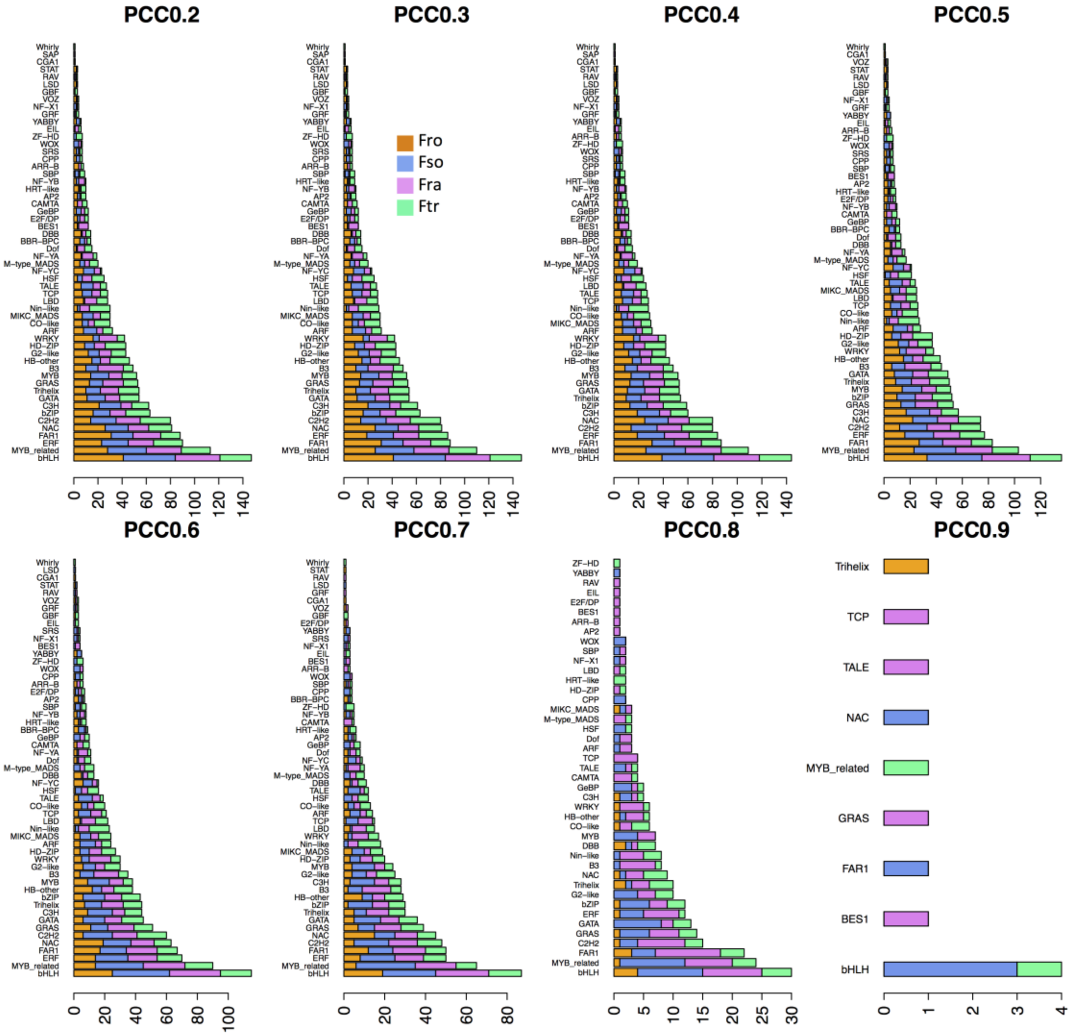


**Figures S7. Distribution of $C_4$TFs for each Flaveria species under different PCCs cutoff**

Bar plots show the TFs number for each TF family (TF pattern) in four Flaveria species under different threshold of PCCs ranging from 0.2 to 0.9. The horizontal axis displays the number of TFs for each TF family.

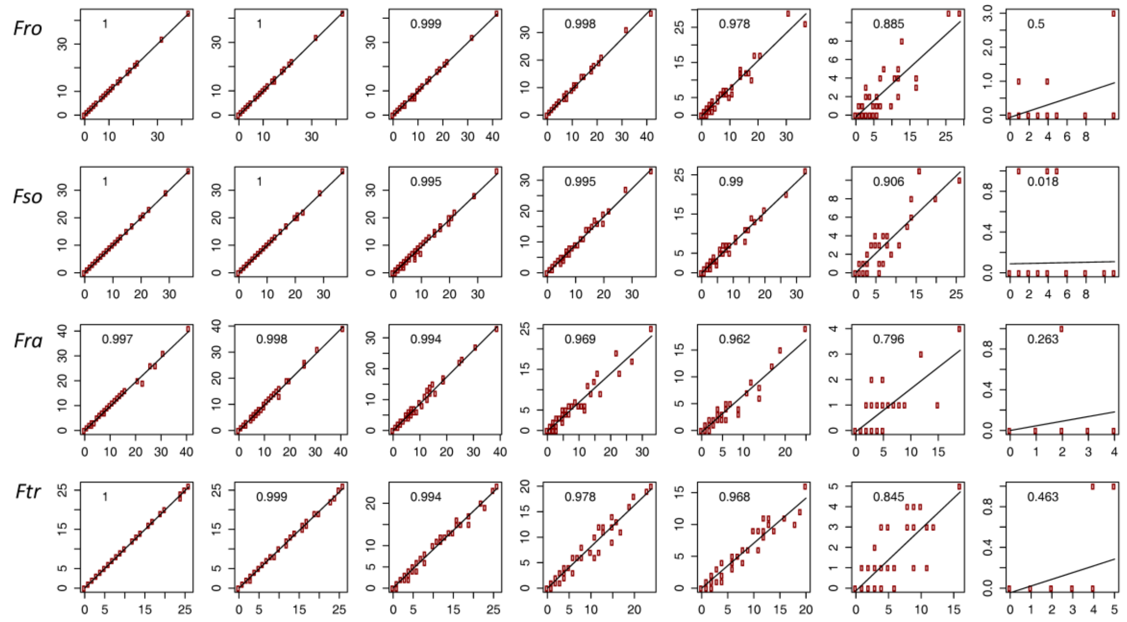


**Figure S8. Determination of the proper PCC for GRN construction by considering the TF architecture**

Each plot shows the correlation (comparison) of the TF patterns for two adjacent PCC pairs. X-axis shows the TF pattern under a high PCC threshold and Y-axis displays that under a low PCC threshold. The number on the top left shows the Pearson correlation between the two TF patterns resulting from the two PCC thresholds. The Flaveria species abbreviations are as shown earlier in Fig. S1

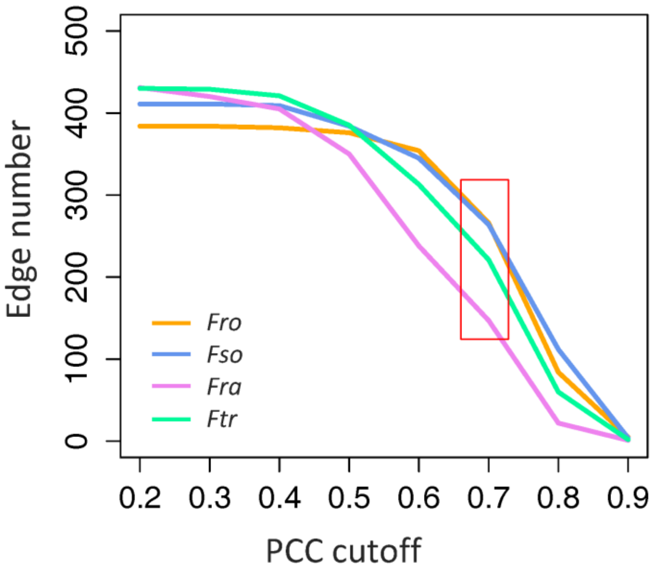


**Figure S9. The edges number of $C_4$GRN under different PCCs cutoff**

The curves show the distribution of $C_4$GRN edges (including $C_4$ core metabolic genes and TFs) in four Flaveria species under a PCC cutoff ranging from 0.2 to 0.9. The number of edges shows an inflexion point at a PCC of 0.7. The abbreviations of Flaveria species are as depicted in Fig. S1.

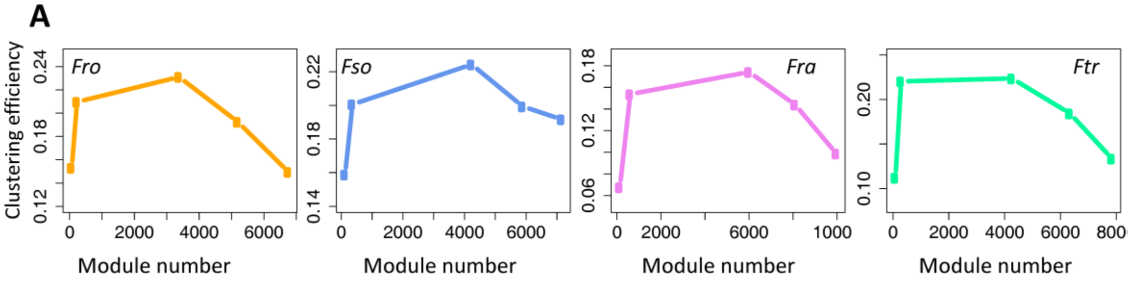


**B**

| **Species** | ***Fro*** | ***Fso*** | ***Fra*** | ***Ftr*** |
|---|---|---|---|---|
| Clustering efficiency | 0.21 | 0.2 | 0.16 | 0.22 |
| # clusters | 222 | 363 | 605 | 295 |
| $C_4$ TFs enriched cluster (s) FDR<0.001 | Cluster No.2 | Cluster No.2 | Cluster No.3 | Cluster No.2 |
| # TFs ($C_4$ genes) in the cluster | 98(2) | 66(5) | 30(2) | 68(3) |
| # genes in the cluster | 3868 | 2469 | 665 | 2107 |

**Figure S10. Statistics of the genome-wide GRN clustering into modules**
(A) The various plots (or curves) show the number of modules and clustering efficiencies at five different thresholds of the parameter "I" of MCL package, *i.e.*, 1.4, 2, 4, 5 and 6. (B) The statistics of genome-wide network clustering under a threshold of "I" to be 2. "I" stands for Inflation. The abbreviations of Flaveria species are as reported in Fig. S1.

**Table S1. Statistics of the transcript assembly and orthologs prediction in four Flaveria species**

| Species | #Transcripts | #ORF[@] | #Transcripts having ortholog in *Fra* ref[&] | % ORF having ortholog in *Fra* ref[&] |
|---|---|---|---|---|
| ***Fro*** **($C_3$)** | 80514 | 80145 | 48092 | 60 |
| ***Fso*** **(I, $C_3$-$C_4$)** | 74137 | 73638 | 43959 | 59.7 |
| ***Fra*** **(II, $C_3$-$C_4$)** | 163967 | 163503 | 116752 | 71.4 |
| ***Ftr*** **($C_4$)** | 78029 | 77683 | 47097 | 60.6 |

[@]: The number of transcripts with open reading frames (ORFs). [&]: Gene reference of *Fra*.

**Table S2. The four conserved TFs co-expressed with PEPC1 in *Fra* and *Ftr***

| ***Fra* ID** | **TF family** | **Ortholog in *A. thaliana*** | **Functional annotation** |
|---|---|---|---|
| Gene03895 | DBB | AT2G21320 | Negative regulation of transcription |
| Gene12432 | NF-YA | AT1G30500 | NF-YA7 |
| Gene17333 | MYB_related | AT1G74840 | Regulation of drought-responsive genes |
| Gene18956 | GRAS | AT1G14920 | Repressor of GA responses |

**Table S3. Primers used in this study for qRT-PCR analysis**

| Primer | Strands | Sequence | Product size |
|---|---|---|---|
| CA1-con | F | GCTCTGACTCTCGAGTTTGC | 162 |
| | R | GCTCCACCTTGAGATGCAGAA | |
| PEPC1-con | F | GCTTACATCATCTCAATGGCCAC | 223 |
| | R | CTGAATCAGAGTACCCAATCATGAC | |
| PEPCk-*Fro-Fso-Fra* | F | GCGGATTTCGGATCGGC | 185 |
| | R | CCGTAAAACGGAGGAACTCCGG | |
| PEPCk-*Ftr* | F | ATCGGTTATTGTCATCAGTTAGGTATTGC | 150 |
| | R | GACAACTCCGGTCATCGTCC | |
| NADP-MDH-con | F | GCTGGGATGATATCCAACCA | 141 |
| | R | GTCCTCAAGTTCCATAGCCAC | |
| PPDK-con | F | GCTGGTCTAGCTGGCAAA | 219 |
| | R | TGCTTCCACATACACGTTCTTG | |
| PPDKRP-*Fro-Fso-Fra* | F | GAGGAGGAAGACGGTTGTCT | 193 |
| | R | CATCGTCAATCCCAGAGAAC | |
| PPDK-RP-*Ftr* | F | CATTCGGTCAACGCTGCC | 179 |
| | R | GCAGCCATGTTTTCATCAGC | |
| NADP-ME4-con | F | GATGATATACAGGGGACAGCTTC | 267 |
| | R | CCAGGGCTTCTTGAAATGC | |
| AlaAT2-con | F | GTGAGTATGCTGTTCGTGGT | 171 |
| | R | ATAAAGCAAGAACCTCTCTGAAGAAAGTA | |
| BASS2-con | F | GCTGTGCTTTGTGGGAATGC | 164 |
| | R | GATATTGTTCTTGAAGAAGACACGTCCAC | |
| DiT1-con | F | GCCGCTGCAGTTCCGGCGAAACAG | 158 |
| | R | GACAGCAATTGCCACGCGTTCTTCGAGACT | |
| DiT2-con | F | GCGACCTCAATCACCGAA | 110 |
| | R | GGCTTCGGGACGACGAA | |
| NHD1-con | F | GTCTTACATCTTATCGGTACCTCCG | 229 |
| | R | GCTTGTTTCATCCACTGAACATAAAG | |
| PPT1-con | F | GCTTTGGTACCTGTTTAACATCTACTT | 182 |
| | R | GGATTGCTAAAAGCTGTGCAC | |
| Actin7-con | F | GTGCTGGATTCTGGAGATGGTG | 162 |
| | R | TTCGGCGGTGGTGGTGA | |

F and R stand for forward and reverse primer sequences, respectively.

**Table S4. Primers used for cloning the promoter sequences of PEPC1**

| Primer | Strands | Sequence | Product size |
|---|---|---|---|
| *Ftr*_PEPC_pr | F | CGTGAGATTGAAAACTAGTCTGG | 1012 |
| | R | ACCTGCGAAAGATTAAATACGAGA | |
| *Fra*_PEPC_pr | F | AGCCCGATCAGCCGAA | 1253 |
| | R | GTCAGCCGCAATCAAACACC | |
| *Fso*_PEPC_pr | F | CATCTAGAACTTGAGGACACGA | 1306 |
| | R | TTTTGCCCTCACATGACAAGC | |
| *Fro*_PEPC_pr | F | ACACACCAGGACACGAG | 1644 |
| | R | CTTTAAGGTCGTATCAGATTTGGTC | |
