## Supplemental File 7 for "Evolution of co-regulatory network of C_4_ metabolic genes and TFs in the genus Flaveria: go anear or away in the intermediate species?"

>*Fro*_PEPC1

ATTGTAAACGAATTCACCTCTTATATATAGAAAAAAAGTCATACCCCTCTTATATACATGAGGTATGACTCTTTT
CTAGTTTGTATAAAAAACAATTATGTATACGTCAGTCGATGGATCTCTCCTCATCAATCCATAAAAAAATGCTTA
TTTTGGTTTCTAATTATTTAATGTTTTTTAATTTCAACTTAGATTCTATTGAAGTTTTACCATGTTATTTTCTTT
TGATCTAAAAACATGCACTCAATTATATACTATAGAGGTAACCTTTGAAATATAGAAGTCAACTCGTTAGAGTTT
TTATTTGACTTAGAAAGTCAACGTTTTATTTCCTATGACAAAATTTGTGGTTCATTGAACCTCTATGGTACGTTT
GATGCCTTTTATTTGTAAATTGTTAAACTTCAACAAATTCGGTTATAAATAAATAGCCGTAAAGCGAACGGCCGT
ATCTTTTACAAAATATAAACTTATCATATTACTCTTGTGATATATATAAAATTAACACAAAAAATAAAAATACAA
ATCAAACCAAACCGACCAACACACTCACTCTAAAATCTACATCTTCTCTCTCTCTCCCTCCCACTCATAACAA
ATTAAAACCTCGTCTCTTTAAACCCATCAAGGCCACCAAGCACTACAACATCTTTATCGCCCATGACCAACGAAC
GCCACCACATATGCTTCCCCGTTCATCTCTCTCGCTCTACCATCATCATCTCTCCATCCACCTACGAATGCCACC
ACCATGTCACACCTCCTATAGCTATCGCCACCTTCTTTGATCTGCTATCCGTCAAGTTCTGCAAATCCAAGTTCA
AGACCCCTAAATCGACTCCCAACAACACCAATCATTCAAGACTCCTAAATCCCAAATCGATCTATCACCACCACC
ACCATCACTAATTGTGATGTTCTAGGTTCAAACAACCCAATCAGATCTAAGGTTTGGTAGATATTAAACAATCCA
GACCATCTTTTTTGTTTCCAAGCCAAAAATCCATTCATCAATCCCTAAATCCAATTTGTGATTTCGGTGGTGGTA
AAAATAGGACAAAGCTCAAATGGTTTTGGAGTTCGTTGATGGTTTGTGGCGATGAATTAGAATGACAAGCAGGGC
CGTCCCCGAGAATCCTCGAAAAATTCTGGGCCTTTGGCGAATAGAAAAAATAGGCGTAAATAATATAAAATTAAT
AATCCGTATGTAAAAGTCTATAATAAAACAATAATTATTATATTACTAAACCAAAATATTGTAGCAAAATAATAT
AATTTACTTATTTAAGCAAATCGTGAGGCTCTTTCAACGTTTTGAGAATGAGTTAAGATTAGGTATATAGTAATG
AGCTAAACTAATTAAAATGTTAATAGACTCTAAAATTTAATATGTTAAAATAATTAAATTGTTATTGGGTTTTTG
TTAATAAGCTATAGTTTTTTTAAATAAAATATACGTACATATACATAATATATATAAAAAATAAAAACTTGGAGT
CCCCTAGACGGCGACCCACCCCGTCCCCCTCAGGGCTACCTATGATGACAAGGGTTCATAAAGATGGCGGTGATG
TTGGTGGACGTGTGTGGAACGTCTGTGGCAATGACTGATGGTCTTTGAAATCAAAGGGGAGGGGTGTGATTGTGG
TGGTTTGGTGGTGTTTGATTGAAGGTGAAGGAGGGTTATAGGAGGTGGTGAGTGATGGTGGTAGTGGCGATGGAG
GAGAAGAGTGAACAATGATGGTTGGCGGTGATGGAATATGACGATACACATACATACACACACACACACACCAAG
AGAGAGAGTTGGGTTGGGGGCTAGAGGGATTATAATTTTTTGTAATAAATTTTAAAAATATGTTAAAAATTTGTG
AAAATAACTAAATTCCATTGAGTGTGATATTGCATCACTGGATTGCGTTAGTTTCTTTAGTAAAATTGAAAGCTA
GTATGAAATTATTAGACTGTTAGCGTAACAAATAAATTTTTTGATTGAATCGTAAATTTTGTCAAACCGTAGAGA
CTATAAGTGTAAACTCTAAAAGTAAATTACATTTAAACTTATTTAATAATTTTAAAGTGTTGTTTAAAAAATAAG
TATATGCTTATGTTTGTTGATAGTTTTTCTTTTGTATTTGTTATTATTTACGAGTAGAACATGAAAAAGACACAC
CAGGACACGAGCATCTGAGTTTTTATTCGAATATTTCTCTTTACTCAACAGAAAAGTAAAACAAATCCATGAAAA
GGATAATTAAGTTATATGTGGACCGTATTGAGACTATATTTTTGTGGTGGAAATCATATGAATTTATGAAAAACT
CATGAAGAAATTGAATTAGAAAGAGGAAATAGAAAGCAAAGTTGGATCTTTCATATCACGAAAAGGCATGAGTTC
TTGCCACTTGACCAAGGAGTGTTCGTAGAGCCGTCTTACTCACTAAAACAAACAAAAAAACAAAAACAAACAAAA
TCTTTCATAAAAGATGAATCGAACACTTTTTCTTTTTTGTCATGATACATATACATATGTATATATATATTGAGA
AATGGAAATTAAGTACAACCGTATTCTCTATATGTAGAGGCGTTAAACGGGTCAAATTTGCGTGTTAAAGACCAA
ATCTGATACGACCTTAAAGATTTTTTATGAATTTGGAAAGTATGCATATACAAAAATTATGTAAGAGATGTGAAC
TTGAAAATGACTTGTGTGTTGATTTTTTTGTCACCTAGAAATCAATATATATATATATATATATATATATATATA
TATATATATATATATATAATTGCACTCGTAACGGAAGATGATTCGTTTTTCATGTTTTAACTTTTTATGTAAAAT
ACCTAATAAGTTTGAGATTTAAAATAAATTAACTAAAGTTAACAACAATAACCAAGTGAACTCATGAACACATCG
TACTGATTTTACACTCACCATTTAGCTAAGTGTGTGTGTGATTCGAGTCAAAAATTCACACATCTAACTATATGC
ATTAGGTGTGAGGGAGCCTAGAAAAATTATATGAGAGGGTTTTATATTAGGAATTTTTATTATCATATAGTTTTC
TTTTGTGTCTATGAGAGCTAATGTGGTTATGATAGGTAAAAAATCACCGAACTTAATATTTGAATATAAGACCTT
TGGTAGGAGAGAATTAAGTATTTACCACTACATGCTTTTGAAGGCTTTCTCATATTTTTCATTGAGAATTACATC
GTTGCTAACTTTTTTGTATAATATCATACATAAATTACTTATATTTTCAAAGACATAATGAGAATCATTTAAGCA
TAAAGTACAATCAATCAATGCATAGAGTTTGATGGCATTGTGACGTGCTTATTTTTATTTACATTTTAATATAAA

TTTTATATGAAATCAACCCGGGTTAAATATAATATATTTTTTAATTTTTTCACGGCATAGTAATTTTCCTAAATT
TCAAATATGTTTAATTTCATTTATTTTTCAATATACATTTTATTTAAAATGAATCACTCGCAATGTGCGACTAAA
ACCCACGGGTCAAGGTATAATGCATTTTTTTTTACACAAACGTAATATTTTCTAACTTTTCCCAACAAAGTGATG
TTCTCCTAAATTTGAATATACCATTATGACATGGTTACATCCACCTAAGTATTAATAATTATTTTATTTAAAATC
AAGCCTTATAAAACGTGGCCGAAACTCACGAATCAAATATAGCTTTCTTCACACTAACGTTTTTATGTTTCAACT
ATACCATTGTAACGTATTTTTTCTACCTACGTCTCTAAATAAATTATATTAAAAAAATGACGCCGCAACACGCGG
AAAAACTACTAGTTGTCTTCTAATTCACAAAAATTCCCAACGAATGATTAGTTGCGTTTGTTATGCATACGAAAG
CGGACGATAATGTCGTTATTATTAAAAAAATACTAAAAGAGTAAAAACTAGAAGAAAAAGACTGATTATCAATTT
ATAATAATATCCACAAAAATATTCC

>*Fso*_PEPC1

ATGACGAGGGTTTACAAGTGGTAGTGGAAGATGAAGGTGATGGTGGTGGACATGAGTGGAACGACAGTGGCAATG
GGCGATGGTCTTTGAAATCAAAGGGGAGGTTTGTGATGGTGGTGGTTTGGTGGTGTTTGATTGAAGGTGAAGGAG
GGTTGTAGGAGGTGGTGAGTGATAGTGGTGGTGGTGGCGATGGAGGAGAAGGGTGAACAATGATGGTTGGTGGTG
ATGGAATATGACGATATACACACATTCAGAGAGAGGGAGAGAGAGGAGAGAGGGAGGAGAGAGAGAGAGAGAGAG
AGAGAGAGAGAAAGAGAGAGAGAGATGGGTTAGGGGCTACTATAATTTTTTGTAATAAATATAAAAATCTGTTAA
TTTTGTGAAAATGACTAAATTCATTGATTGTGAAACTGCATCAGTGGATTGGTTTAGTACTTTGGTGAAATTGAA
AACTAGTTTGAAACTATTAGATTGTTAGCGTAATAAATAAATTATTTGACTGAATCATAATTTTCATCAAACCGT
AAAAACCACAAATGTAATTTACTATAAAAATAAAATAAATTCAAACTTGTATTCGAATTAAACCACTACTATTTT
ACTGCTATGAAGAATATTTAATAATTTTAAAGTGTTGTTTCAAAAATAACTATATGCTAATGTTTGTTGGTAGTT
TTTCTTTTGTATTTGTTATTATTTCATCTAGAACTTGAGGACACGAGCATCTGAGTTTTTATTCGAATATTTCTG
TTTAATCAATAGAAAAGTAAAACAAATCCATGAAAACGATAATGTGGACCATATTGAGACTATATTTTTGTGGTT
GAAATGATATGATTTTATGAAAAACTCGTGAAAAAATTGAATTGGAAAGAGGAAATAGAAAGCAAAGTTGGATCT
TTCATATCACGAAAAGGCATGAGTTCTTGCCACTTGACCAAGGAGTGTTCGTAGAGCCGTACTTACTCACTAAAA
CAAACAAAAAACTAAAACAAACAAAATCTTTCATAAAAGATGAATCGAACACTTTCTTTTTTGTCATGATACATA
TACATATGTATGTGTGGGTGAATATATGCATATATATTGAAAAATGGAAATTAAGTACAACGGTATTCTCTATAC
ATAGAGGTATTAGCGGGTCAAATTTGCGTGTTAAAGACCAAATCTGATACGACCTTGAAGATTTTCTATGAATTT
GGAAAGTATGCATATACAAAAATTATGTAAGAGATGTGAACTTGAAAACGACTTGTGTGTTGATTTTTTTGTCAC
CAAAAAATCAAATACCTGTAAAATAGTTGCACTCGTTACGGAAGATGATTTGTTTATTAATGTTTTAACTTTTTA
CATAAAATACCTAATAAGTTTGATATTTAAAATAAAATAACTCTTAAAGTTTAACCTGAATAACTAGGTGAACTC
ATGAGCACATCGTATTGACTTTACACTCACCCATTTAGCTAAGTGTGTTTGTGATACGACCCTATTTCGTATTAT
GTCAAAAATTCACACATCTAGCTGTATGCGTTAGCCATTAGGTGTGAGAGAGCCCGGGAAAGTTTTATGGGAGGG
TTGTATGTCCGGAAACTTTATGATCATATAGTTTTTTTTTTTTGGTGTGTGCGTGTGTGTCTATGAGAGCTAATG
AGGTTACCATAGATAAAAAAAACCACTGAAATCCATATTTGAACATAATACCTTTGGTAGAAGAGAATTAAGTAT
TTATGACTACATACTTTTGAAGGCTTTCTCGTATTTTTCACAGAAAACTACGTTGTTGCTAACTTTATTGTATAA
TATATGGCACATAAATTATATATAGAGTAAATTACACTTTTCGTTCTTTAGGTTTATAGCAAATTTCAGGTTTCG
TCCTTAAACTAAGAAAATTACAGTAAACTTCCTTTACATTTAAAAAAGGTTGCACATTACATCATTCTCACTAAC
GACTGTCTATTTTAGACGTTAAATAAGTGCATGTGCTTGTCATGTGAGGGCAAAATGGTCATTTTATATTTTATA
TTTGAAATTAAAATTAAACAAATAAAAAAAAAATACTTTTGCCTTTTTCTTTCTCTTAACCATGAAATCTGAACC
ACACCACAGAAACAACCCTGCAACTCAAATCACTACAACCCTCTAAAACTCCGATCAATCTACCCCTTCAAAATT
AAGGCCACCATCACCCCCTGCATCTTCGGCAACAACCACCATTGTCTATTACTCAATCTCACATCTCTCTCTCTC
GCGCTACCATTTCTGGCAAAGGTGGTGGCGATTAGATCGGCTTTTGGCTCCGAGGTCCGGTTGCTAGATGATTTG
TTAGATTTGTTCTTGCTTCTGAGCTGCGGTTGTGGCTGGAGGTCACGGTGGTCGGCGGTGGTGACACCAGCACTG
TAAATGCAACATGATTGACTGATTAAGGGGCTTAGTGAGGTTGTGATAGATGTATTGTGTTCTCCATGTGTAGAA
ATAAACTGTTGAGGGTGGGCGACGAGTGGTGGCCGACGGAGGAGAGGCGGTGGTGGTCGTTGAAGGATTAGGTTA
GGAGAGATATGTATGAAGCTTTTTTGTATGGTGAAATTGGGATTTGGGGTTTATCAAGAAACCCAAATTGAAATG
TGAGATGAATCAAAATACCTCAGATCGAAAAACTTCAAATTGGGTGTGGATTGAAGAAAAAATTAAAGAATCATG

TTTAAAATCAATAAAAAAAAGGGTTTCAAGAGTTTTACTCAAAACTCAGACATCAAAAGATGAAATCAAAAAATC
CTAACTCAAAAGGTTTAAGAATGAGATAAAATTTCATGTTAAAAAGTGAGATGATAATATACAAAACAAGAAACC
ATCACTGGGATTTCAAAAAATCTTTCATATTTATAAGGGATAAAAAAAACCAAATGAAAGAAAATTTTCACATTG
AATGTGTGAGGGTGAGATATCAGGTTAGCAGTGGGGTTGGGTGGGGTGATTGAGGATGTGGCCGGAGGTTGTTGG
AGACGGTGGTGGCCGTTGGCGGTTGCTTTTATGTAGAGAGAGAGAGAGATAAAGAGTTAAAGTAGGTGGTGAAAT
AAGTTCAATAATTTAGTTAATATAATATTTTTTATTAATAAGGGTAAAAATACTAATTTATCCTCACATGCAGGT
CACATGATCTGATTTAACGGACAAAATTAACGTCCGTTAGAGAAAAGTACGAAACGTGCAAGCTTTTTTAAACGT
AAAGGATGTTTACTGTAATTTTCTTAGTTTAAGGACGAAAACTGAAATTTGCTATAAACTTAAAGGACGAAAAAT
GTAATTTACTCTTATATATATTACTAACTATCAAATTCTTTATTCTTTATGGTTTGTACTGTAATCTTTAGTGGG
TTTAGGAATTTTTTTACACCGCAATACTTCTACATTTTTTATAACTTAACCCTTATAATTTTTTTACTTTAAACT
TTGATCCCATTTACTCTTTATATTTTACAAAATTTTCGTTTTACGTTTCAGTCTAAATATTACGAGTTAACATGC
CGGAACGTGCGCGTGTGGCTTCAATGTTTTTTACACATATTTTTTCCCGTTTAACCGCTCCGTCACAAAGCACGA
GTCATAGATTGACTTAGTTATTATTTATAATATTTTACATTTCTCGTGCTTGTTTCAACGTTTTTATGTCTACTT
TTTAGGTTGGTTTATTTTTTTGTTGTTTTTTATAATATGAGTACTTCCGGTTTTGATAATCGATGACGGTAATGT
AATATTGATGTTAAATGACATGGTTTTATACCCCCCTATAACGGTGGGGTTTAATACCAGTTGTTTTCTAATTCA
CAAAACTCTCGACGAATGATTAGTTGCGTTAGTTATGCATACGAACGCGGACGATCATGTCGTTATTATTAAAAA
AATATCAAAAGAGTAAAAAATAGAGGAAAAAGACTGATTATTAATATAATAATAATATCCACAAAAATATTCCAA
TAATTCAACCCAGAGTTTGCTCTTT
>*Fra*_PEPC1
ACCTTGGAAACTCACCAACTATTTATAGTTGACCTTTTATTTTTATCATGTTTCAGGAAAACCCCCAAGTTTTGA
AGTGAACTTATGTTGCTGGCATATTCTAGGACTTAGACCTAGTTAGTAAACAAAATGATTATGTTTGTTATTTCA
TGTTGAAACTTATTTCATTATTGAGACTTATGCTTGTAATGTAATTTTGAAAGCTTATGAATAATATATTTGGGA
TTTAATAATAGCGTTATGTGGCTGTGAACAATCTTAACCCTCACACCTCGACGTTTCCGCCGTCAAGGTTGGGGT
GTGACAGATCTTACTGAAACTTGGCTATTTGATTCAGTATACGAATGAGTCAAAAACCGTATATTTTCTAATCAA
CTAGAATCATTAACAATCTTTAATCAACTACTAAATCAAAATCGAAACCACCACACCATCACCACACCACCACCA
CAACCACCACCATCTTCGTCACCTCCAACAATAACAATCAATCTTCCACCCAAAAACCTACTGCAACAAACCACA
CCATTTCTGATAAATCGAAGCTCCATAGATCCCCCTCCATCATCCACCATCTCAAATCCTTCACTACCCACCACC
ACCACCAAGATCTACCACCACCACCTACCACTTTCGATTCCAATCCGGAGGGAAGGTTAACATAACGAAGTTCGG
ATGAAGGAGCGCCGGACGGCGGCGACGGAGACTGCCTGCCAGTCGCGAGGGGGCGACTGCCTGCGGGTCGCGAGG
GGGTCGTCGACGGGTGTGCGGGGGTGAACTACGGTCGCCCCTTTTCTTGGTCTTGGGGAGGAATCGCGCAGAGGA
GGAATCGCGCAGGGGAGGTGCTCGGGGGTGGGTCTTCCGGCTGCGCAAGGGGGGTGAGTGGGGTGGATCGGCCGG
CTGCGCAAAGGGGGGTGGGTGGGGTGGATCTGCTGTGTTTTTGATGAAGATGGATGGGTCAGCATTTTTTATTTT
CAGGTTTGGGTCAGCTGCAAAGGATCCGGTCAGCTGATACCAATTGCATATCAAACGGGCTGACCCGCTGACCGA
GTCAGCCAGTCAGTCGGGTCAGCCCGATCAGCCGAAATCAAACACACCCTAAGTTCCATTTACTGTAAAGTTGCA
TGAGTGGATTGGTTTAGTGTCTTTCGTGAAATTGAAAACTAGTCTGAAATTATTAGACTAAATCATAATTTTCAT
CAAACAAATAAAGACTATAAGTGTAATTGACAATAAAAATAAAATAAATTCAAACTTGTATTCGAATTAAACCAC
TACGTATTATTTCACTGACATGACTAATATTTAATAATTTTAAAGTGTTGTTTTAAAACTAACTATAAGCTTATG
TTTGTTGATAGTTTTTCTTTTTGCATTTGTTATTTTTTTACGTCTAGAACATTAAAAAGACACCAGGACACGAGC
ATCTGAGTTTTTATTCGAATATTTCTCTTTACTCAATAGAAAAGCAAAACAAAATCCATGAAAAGGATAATGAAC
TTATACGTGGACCATATTGAGACTATATTTTTGTGGTTGAAATCATGTGAATTTATGAAAAACTTGTGAAAAAAT
TAAATTGGAAAGAGGAAATAGAAAGCAAAATTGGATCTTTCATATCACGAAAAGGCATGAATTCTTGCCACTTGA
CCAAGGAGTGTTCGTAGAGCCGTACTTACTCACTAAAACAAACAAAAAAAAAAAAGTAAAACAAACAAAAAACTGA
AACAAACAAAAAACTAAAACAAACAAAAAACTGAAACAAACAAAATCTTCCATAAAAGATGAATCGAACAATTTT
CTTTTTGTCATGATACATATACATGTGTGTGTGAATATTTGTATATATATTGAGAAATGGAAATTAAGTACAACG
ATATTCTCTGTACATAGAGGCGTTAACGTGTCAAATTTGCATGTTAAAGACCATATCTGATAGAACCTTGAAGAT
TTTCTATGAATTTGGAAATTATGCATACACAAAAATTATGTAAGAGATGTGAACTTGAAAACGACTTGTATGTTG

ATTTTTTTGTCACCAAGAAATCAAATACCTATAAAATAATTGCACTTGTTATGGAAGATGATTTGTTTATTTATG
TATTAACTTTTTACATAAAATACCTAGTAAGTTTTACTTTTAAAAATAAAATTACTCTTAAATTTAACCAGAATA
ACCAGGTGAACTCAAAAACACATCATACTAACTTCACACTCACCCATTTAGCTAAAGGGGTGTTTGATTGCGGCT
GACCCGGCTGACCCGGCTGACTGGCTGACCCGCTCAGCGGGTCAGCCCGTTTGATATGCATTTTGTTTCCTGCTG
ACCGGGTCTAAGTAGCTGACCCAGACCTGAAATTAAACAAAGCTGACCCAGACCACCGAGTGACCACCGACGGCC
ACCACCCTACCCACCCTCCTTGCGCGACCACCGAACAACCACCGACGGCGGACCCCCTGCGCGACCAGCCCCCTG
CGCGACAACCCCCCCCCCCTGCGCGACCAGCCCCCTGCGCTGCTTGACCAGCCCCTTGCGTGACGAACTCCCTGC
TCTTCGACCCCCCCTGCGCGACGACCTCCCTGCTCGACGACCCCCCCTCCGGCGACGGCGTTTCCGGCGGAGATT
GTGGTGGTGGTGGTGGTGGATTTGGGGAGGAGTAAGGAGGTGGGTTTGATGAGGGGTTTTGAGGGCTTGTTGGAG
GTCATGCATTTGGCGTTGAGCATGGCCATGGTGGTGGAGAGTGTGGCTGCCATCGGTGGTTGTTGTGGTGGTGGT
TGTTGTTGTGGTGGTTGATGTGGATGAGAGGTGTGTGTGAATGTGCATACACGAAGTGAGAGTCAATGGGGTTAA
TGTAATGGAGATTGTGGTGTTATCCATTGGATTTCACATTTTTTCTAAGTAAATTTCATATTCTGTTAATGTGGT
TTGATTCAAATGATGTTTTTTCTTCTAAAGGTTTTAAAAATTATGATTTTACTTGTAGTTAATAATTTGTCATAA
TTAACTTCGATTGACATCAATTGACCAAACGGGACAATATGGTTATTTTTCTTATTCAGCAAAGTCAGTCAGATG
CTAAATCAAACAGCCAAGTTTTAGTCAGATGCAGACCCAGTCAGAACTTCTTACTGGGTCAGCTTCTGACTCAGT
CAGCAGTAATCAAACAACCCCTAAGTGTGTTTGTGATATGACCATATTTCGTATTATGTCAAAAATTCACAAATC
TAGCTGTATGCATTAGGTGTGAGATAGCCCGAGAAAGTTCTATGGGAGGGTTGTATGTCCAGAAACTTTATGATT
ATATAGGTTTTGTTTTATTTTGTTTTGTTTTTTTTTTGTTTTGTTTTTTTGTTTTGTTTTTTTGTTTTTTTGTTT
TTATTTTGTGTGTGTATGAAAGCTAGTGAGGGTACCACCGATAAAAATACCACTGAAATCGATATTGAATATAAA
AATACCTTTGTTTGAAGAGAATTAAGTATTTATGACTACATACGTTCTCGTATTTTTCACTGAAAATTACATCGT
TGCTAACTTTATTGTATATATCTCGTACATAAATTATATATTACTAATTATTAAATTCTTAATCTTTTATGGTTT
GTACTTTAATCTTTAGTGGGTTTAGGATTTTTTTACACCCCAATGCTTCTAAATTATTACAACTTAACCCTTATA
ATTTTTTATATTCAACTTTGATCCCATATACTTTTTATATTTTATAAAATTTTCATTTTACTTTTCAGTCTAAAT
TTTACGAGTTAACACGTCGCAACGTGCGCGTATGACTTTAATGTTTTTACGCATATTTTCCCATTTAACGTTTCC
GTCACAATGCGCGAGATGGATTGACTTAGCTATTATTTTTTTAATATTTTACATTTCTCGTGCGTGGTTCAACGT
TTTTATGTATAATTTTCAGGTTGGTTTATTTTTTTTATTGTACTTTATAATGCGAGTCTTTCCGGTGTTAGTGGT
CGATGATGTTAAATTACATCGTTTT
>*Ftr*_PEPC1
ATTTATGATCTTCGGCCCAGTACCCACTTTGTGCATACGTTAGAGGTGTAGTATACGGTTGGTATCCTTGTCACA
TGCCCCAGTGCTGGTTCCATGATATGTGCGTTATGAAAGTCATTTATGATGTTCGGCCCAGTACCCGCTTTGTTC
ATACGTTTGTGGTATAGCTGATTACATTAATCTGAAAAGAGTTAAAGAGATAAAGAAAGAAAGTGATGAACAGTA
AAGGAAGATAATAATGTTATTGTCTATGTGTACGTACGATGTATGATTATGTACGAAAGTATAGTATGATTTGAA
AGAAAGTAAAGAATGGTTTTAAATGAATGTAAAGCATGTTTTATTTCAAAAATAGAACTCACTCAGCAATGCTGA
CTTTTATGTGTTTCCAGTTGGCAGGTAAAGATTTTTGAGGATGGCACGGGGCGGCTACTCGGCTTTTTTTTATGG
CATCCATTTTTATAAAGCTTTTGTGGTGTAATGCAACAAGTATTTTGTAATTCAATAATGATGTATTATGCTGCA
AAAGCTTTTATAAAGAAATAAAATGTTTGCTTTAATGTTTCCGCTGGAATGTTTCTTTAAGTTAATGAAAATATA
AGTTTTATGTGTAATGATTGATCAGTTGTAAGCGTCTCGGGAACGGAGCCTTACATCCCTCACATTCTCTCTCTT
TCCCTACCACCCTTTCCTTCTCTCTCTAGACCAAATTAAAACTTTGTCTCCTTAAAGGCACCCACAACCACCAAA
GCACCGCCACATCTTCATTGTGCACAACCAACAACACACACGACATATGCATCCTCGGTCATCTATTTCACTCTA
ACATTATCATCTCCATCCACCTACGAATGTTGTCACCAACTCACACCTCCCACAACTATCTCCACCTTTGAAACT
ATATTCATCAAGTTCTACAAATCCAATTTCAAGACCCCTAAATCTATTCCCAATAATACTAATATTAGAGACCGT
TAAATCCCAAATCGGTCCATCACCACCACTCCTTTGTTCTAGGTTCAAATAACCTAATAGATCTATGGTGGTAGA
TATTAAACAATTCAATCAAATCAAAGTAAAAATCTAGACCATCTCCTTTGTTTCAAAGCCGAAAATCCATTCATC
AATCCTTAAATCCAATTTGTAAGTCTAGCGGTGGTATAAATCGGACAAAAACCAAATGGTTTGGGAGTTCATCGG
TGGCATGTGATGATGAACTAGAATGACATGTAGGATCGTCTTGATAATCCAGAATAATTTTAGGCCCTAAGCGAG
TAAAAAAGACACCTATAATATAAAATTAATAACAATATGTAAAAATTATCAAATACAATAACAATTTTTAAAGAA

AAAATATTATATAAAAACAATATAATTTACTTATTTAAGTAAATTGTGAGGCTCTCCTCGCATTTAGGAATGAGT
TAAGATTGGATAAGTAGTTGTGGGCTAAACTAATTAAGATGTCAATTAGCCATAACGGTTAATATGTTAAAATAA
TTAAAATGTTATTGGGTTTTGTTAGTGGCCTAGAGTTTTTTTAAAATAAAGTATACATACATATATAATATAAAA
ATTTTAAAAACTCTGGGCCTCATAGGGAGATTGGGTGCCCTAGGCAGTCGCCTACCCGTCCCTCGTCAGGGCCGC
CTATGATGACAAGGGTTCACAAGTGGTTGTGATGATGGTGGTTTGGTGGTGTTTGATTGAAGGTGAAGGAGAATT
GTAGGAGGTGGTGAGTGATAGTGGTTGTAAGTGATAATGGTGGTGAGTGATAGCGGTGGTTGTGGTGGTGGTGAT
GGAGGAGAAGGTGAACAATGATGGTTAGTTGTGATGACATATGATGATATACACACATTCAAAAAAAGACAGGAA
GAGAGGGAGGGAGAGATGAGGTAGGGGCTATTGGATTATATTTTTATGTAATAAATATAAAAATATATTAAAATT
TTATGAAATGACTAAGTTTGATTTAGTGAGAAGTTACATGAGTGGATAGGTTTGGTGTTTTCGTGAGATTGAAAA
CTAGTCTGGAATTATTAGACTGTTAATGTAATAATAAATTATTTGACTGAATCATAATTACATCAAAAAATAAAG
GCTAGAAGTGTAATTGATTACAAAAATAAAATAAGTTCAAACTTGTATTCGAATTAAGCCACTATGTACTATTTC
ACTTACATGACTAATATTTAATAATTCTAAAGTGTGGTTTTAAAAATAACTATAAGCTTATGTTTGTTGGTAGTT
TTTCTTTTTGCATTTGTTATTATTGATATCTAGAACATGAAAAAAGGACTCACCAGGACAGGAGTATTGCATCTA
TGTTTTTATTCGAATATTTCTCGTTACACAATAGAAAAACAAAACAAATCCACGGAAAGGATAATGAGCTTATAC
GTGGACAATATTGAGACTATATTTCTATGGTTGAAATCATAAAACAAACAAAAAACTAAAACAAACAAAAAACTG
AAACAAACAAAATCCTCCATAAAAAGATGAATCGAACATTTTCTTTTTGTCATGATACATATATATATATATATA
TATATTATATGTATATTATATATATATGTATATATATATACACACACACACAAATATATATATATATATAT
ATATATATGTATGTATGTATATATGTATATATATATGTGTGTGTGTGTGAATATGTTGCATATATATTGAGAA
ATGGAAATTAAGTACAATAATATTCTCTACATAGAGGCGTTAATGCGTCAATTTTGTGTGTTAAAGACATAATCT
AATACAACATTGAAGATTTTCTATGAATTTGGAAAGTATGTATACACAAAAATTATGTAAGAGATAAAAGACTTG
TGTGGAAATCAAATACCTATAAAATAATTGCAATTGTTACGAAAGATGATTTGTTTATTTATGTATTAACTTTTA
CATAAAATACCTACTAAGTTTGATTTTTAAAATAAAATTACTCTTAATTTTAGCAGAATAACCAGGTAAACTCAT
AAACACATGGTACTGACTTTACACTCACCCATTAGCTAAGTGTGTTTGTGATACAACCCTATTTCGTATATGTAT
AAAACTCACAAATCTAGTTGTATGCATTAGGTGTGAGATAGCCCCAGAAAGTTTTATGAGAGGGTTGTATGTCCA
AAAACTTTATGATCATATAGGTTTTGTTTTTTTGTTTTTTTGTTTTTTTGTTTTTTTGTTTTTTTTTGTTTTTTT
TGTTTTTTTGTTTTGTTTTTTTTTGTTTTGTTTTTTTTGTTTTTTTTGTTTTTTTTGTTTTTTTGTTTTTGTTTT
TGCGTGTGCGTGTATGAGAGCTAGTGAGGGTACCACCGATAAAAATGCCACTGAAATCGATATTGAATATAACGA
CCTTTGCAAGAAGAGAATTAAGTATTTATGAGTACATACTTTTGAAGGCTTTCTCGTATTTAATCTTTCGCAGGT
TTAAAAATATTAATTATATATTACACCCCAATGCTTTTAAATCTTTACAACTTAATCCTTATAATTTTTCATTTT
CAACTTTGATCCCATATACTTTTTTATATTTTATAAAATTTTTATCTTACTTTTCAGTCTAAATTTTACGAGTTA
ACAAGCGGCAACGTGCGCGTGTGGCTTCAATGTTTCTACGCATATTTTTCCATTTGACGGCCCCGTCACAACGCA
CAAGTCATAGATAGACCTAGCTATTATTTTTTTTAATAATATTTTACGTTTGTCATGGGTGATTCAACGTTTTTA
TGCATAATTTTCATGTTGATTTATTTATTTTGTTGTACTTTATAATGCGAGTATTTCCGGTGTTAATGATGGATG
ATGTTAAATGACATCGTTTTAATACTAATTGTTTTTTAATTTACAAAACTCTCAACAAATGATTAGTTGGGTTAG
TTATTCATAGGAAAGCGGACGAGCATGTCGTTATAATTAAAAAAATATCAAAAGAGTAAACAAAAAAGGA
